# Retinoid X receptor alpha is a spatiotemporally-specific therapeutic target for doxorubicin-induced cardiomyopathy in adult zebrafish

**DOI:** 10.1101/490706

**Authors:** Xiao Ma, Yonghe Ding, Hong Zhang, Qi Qiu, Alexey V. Dvornikov, Maengjo Kim, Yong Wang, Matthew Lowerison, Joerg Herrmann, Stephen C. Ekker, Tzung K. Hsiai, Xueying Lin, Xiaolei Xu

## Abstract

While the genetic suppressor screen is efficient in suggesting therapeutic genes, this strategy has yet to be successful for cardiomyopathies in vertebrates. To develop such a strategy, we recently established a mutagenesis screen platform in zebrafish for systematic discovery of genetic modifiers of doxorubicin-induced cardiomyopathy (DIC). Here, we further revealed both molecular and cellular insights of the first salutary modifier emerged from the screen, i.e. *gene-breaking transposon* (*GBT*) *0419* that affects the *retinoid X receptor alpha a* (*rxraa*) gene. First, by rescuing the mutation in tissue-specific manner with multiple Cre-loxP systems, we demonstrated that the endothelial, but not myocardial or epicardial, function of *rxraa* is primary to this cardioprotective effects. Next, we showed that the *rxraa*-associated salutary effects on DIC were conferred partially by the activation of retinoid acid (RA) signaling. Finally, we identified isotretinoin and bexarotene, 2 US Food and Drug Administration-approved RXRA agonists that are effective in treating adult zebrafish DIC when administered during the early, but not the late, phase of DIC progression. Collectively, we provided the first *in vivo* genetic evidence in supporting *RXRA* as the therapeutic target for DIC, and uncovered a previously unrecognized spatiotemporally-restricted mechanism for this gene-based therapeutic strategy. Our study also justified that searching salutary modifiers via zebrafish mutagenesis screen can be effective in discovering new therapeutic targets for cardiomyopathies.

## Introduction

When subjected to chronic biomechanical stresses, a heart undergoes pathological structural remodeling, known as cardiomyopathy, that can lead to heart failure (1), a devastating disease that affects about 2% of the adult population worldwide (2). Since the discovery of *Myosin Heavy Chain 7* (*MYH7*) as the first causative gene of inherited hypertrophic cardiomyopathy (3, 4), nearly 100 genes have been identified that either cause or predispose the development of cardiomyopathies (5), launching the era of molecular genetic studies to discover therapeutic strategies. Traditionally, the search of a therapeutic gene target needs to be guided by meticulous mechanistic studies on corresponding disease model, which can be a prolonged and inefficient process (6). In contrast, the phenotype-based genetic screening can be a powerful method to rapidly reveal new gene targets of therapeutic potential, as evidenced by those genetic suppressor screens in lower animal models (7-12). However, conducting genetic suppressor screen can be extremely difficult in vertebrates because of the challenge of colony management efforts (13-15). To overcome this bottleneck, we turned our attention to the experimentally accessible zebrafish model, and have successfully established an insertional mutagenesis screen-based platform for discovering genetic modifiers of cardiomyopathies (16-18). While most of the genes identified to date had been linked to deleterious modifications, some appeared to be salutary genes (18), prompting a new strategy that is equal to a suppressor screen for cardiomyopathies. However, because the screen was conducted only based on fish survival (18), it is critical to further prove that the salutary modifying effect is a cardiac event, and that the affected gene is indeed a therapeutic target.

*Gene-break transposon 0419* (*GBT0419*) is the first salutary mutant identified from our pilot screening of 44 zebrafish insertional cardiac (ZIC) lines, based on an adult zebrafish doxorubicin-induced cardiomyopathy (DIC) model (16, 18, 19). This insertional mutant is due to the integration of *pGBT-RP2* (RP2) gene-breaking transposon element (20), which includes both a protein-trap and a polyadenylation signal (polyA) trap, at the *retinoid X receptor alpha a* (*rxraa*) locus, a zebrafish ortholog of human *Retinoid X Receptor Alpha* (*RXRA*). In a cell nucleus, retinoid X receptors have been known to bind preferentially to the 9-cis form of retinoic acid (RA) and transcriptionally regulate a spectrum of biological processes such as embryogenesis, hemostasis, and xenoprotection (21). The identification of *GBT0419* allele as a genetic modifier from this unbiased screen indicated that RA signaling could be important in the pathogenesis of DIC; a view also independently supported by the identification of *Retinoic Acid Receptor Gamma* (*RARG*) as a DIC susceptibility gene reported from a recent genome-wide association study (22). The fact that *GBT0419* mutant exerts salutary modifying effects additionally suggested the hypothesis that RA signaling could be a therapeutic pathway for treating DIC.

In fact, the RA-associated cardioprotection has separately been substantiated by several pieces of the pharmacological evidence. The first is from a series of *in vitro* studies on the cultured neonatal rat ventricular cardiomyocytes, in which RA had an attenuating effect on multiple types of chemical-induced cell hypertrophy (23-27). More recent *in vivo* studies have shown that administration of RA signaling activators can exert therapeutic effects on a rat model of aortic-banding-induced heart failure (28), a rat model of diabetic-induced cardiomyopathy (29, 30), and a mice model of ischemic-induced cardiomyopathy (31). However, *in vivo* supporting genetic evidence of RA-based therapies on heart diseases is still lacking to this end.

Functioning as one of the primary RA receptors, RXRA plays a vital role in the cardiac tissue as outlined by earlier genetic studies. In mice, the *Rxra* null mutant is lethal, starting from embryonic day 14.5, because of heart failure as a consequence of thinning of the myocardial compact zone (32). Subsequently, a cell non-autonomous mechanism was further elucidated on the functions of *Rxra* during the morphogenesis of cardiac chambers (33, 34). However, an unexpected observation was that mice with cardiomyocyte-specific overexpression of *Rxra* showed phenotypes resembling dilated cardiomyopathy in adulthood (35, 36); this finding dampened any further pursuits on the therapeutic potentials of *RXRA*, underscoring the arduous nature of the traditional approaches to search for a therapeutic target.

In this study, we conducted detailed characterization using adult zebrafish to decipher how *RXRA* can be manipulated in a therapeutic manner for treating DIC *in vivo*. At the molecular level, we found that the salutary effects noted in *GBT0419* could be achieved with either *rxraa* overexpression or RA signaling activation. At the cellular level, we showed that endothelial, but not myocardial or epicardial, expression of *rxraa* is substantial for these therapeutic effects. We identified 2 US Food and Drug Administration (FDA)-approved RXRA agonists that can be repurposed for the prevention of DIC *in vivo* while administered during early phase of DIC. Together, our data demonstrate the power of zebrafish model for discovering spatiotemporal-specific mechanism that informs future translation of the *RXRA*-based therapy for patients with DIC. Importantly, this study of the first salutary modifier validated the concept that zebrafish modifier screen approach is able to discover new therapeutic targets for cardiomyopathies.

## Results

### Endogenous *rxraa* is expressed in both myocardial and endothelial cells

To investigate if the salutary modifying effects of *GBT0419* are of cardiac nature, we first defined the cardiac expression of *rxraa*. Zebrafish have 2 annotated orthologs for human *RXRA*, termed *rxraa* and *rxrab* (37). Both orthologs code for proteins that are >80% identical to mouse Rxra and human RXRA (Supplemental Figure 1), with highly conserved functional domains (PDB #A2T929, PDB#Q90415). By re-examining our published RNA sequencing datasets (GEO, #GSE85416) (38, 39), we observed that the *rxraa* transcript level is 3.4 times higher than *rxrab* transcript in embryonic zebrafish hearts (0.85±0.03 vs 0.25±0.02 reads per kilobase per million mapped reads [RPKM]) and 4.0 times higher in adult zebrafish hearts (1.17±0.07 vs 0.29±0.11 RPKM) (Supplemental Table 1), suggesting that *rxraa* could be the dominant *RXRA* ortholog in a zebrafish heart. We confirmed *rxraa* expression in adult zebrafish hearts using semi-quantitative PCR (Supplemental Figure 2A).

In GBT mutants, a monomeric red fluorescent protein (mRFP) gene in the RP2 vector is fused in-frame with the N-terminal part of the affected gene (20), which reports the expression pattern of the trapped gene as a chimeric protein (17, 40, 41). Because *GBT0419* harbors the RP2 element in the 1^st^ intron of *rxraa* (Figure 1A), we documented mRFP expression pattern in *GBT0419*, and crossed it with *Tg(fli1a:EGFP)* (42) to label endothelial cells and *Tg(ttn:actn-EGFP)* (17) to label cardiomyocytes, both with enhanced green fluorescent protein (EGFP), respectively. By imaging dissected zebrafish hearts at 6 days post fertilization (dpf), we noted colocalization of mRFP^+^ cells (representing endogenous *rxraa* expression) with EGFP^+^ cells in both transgenic backgrounds (Figure 1B, 1C; Supplemental Figure 2B), suggesting the endogenous *rxraa* expression in both myocardial and endothelial cells. This observation is consistent with a previous study that used semi-quantitative PCR to assess RA-receptors expression in purified zebrafish cardiac cells (separated by fluorescence activated cell sorting) (43).

**Figure 1.**
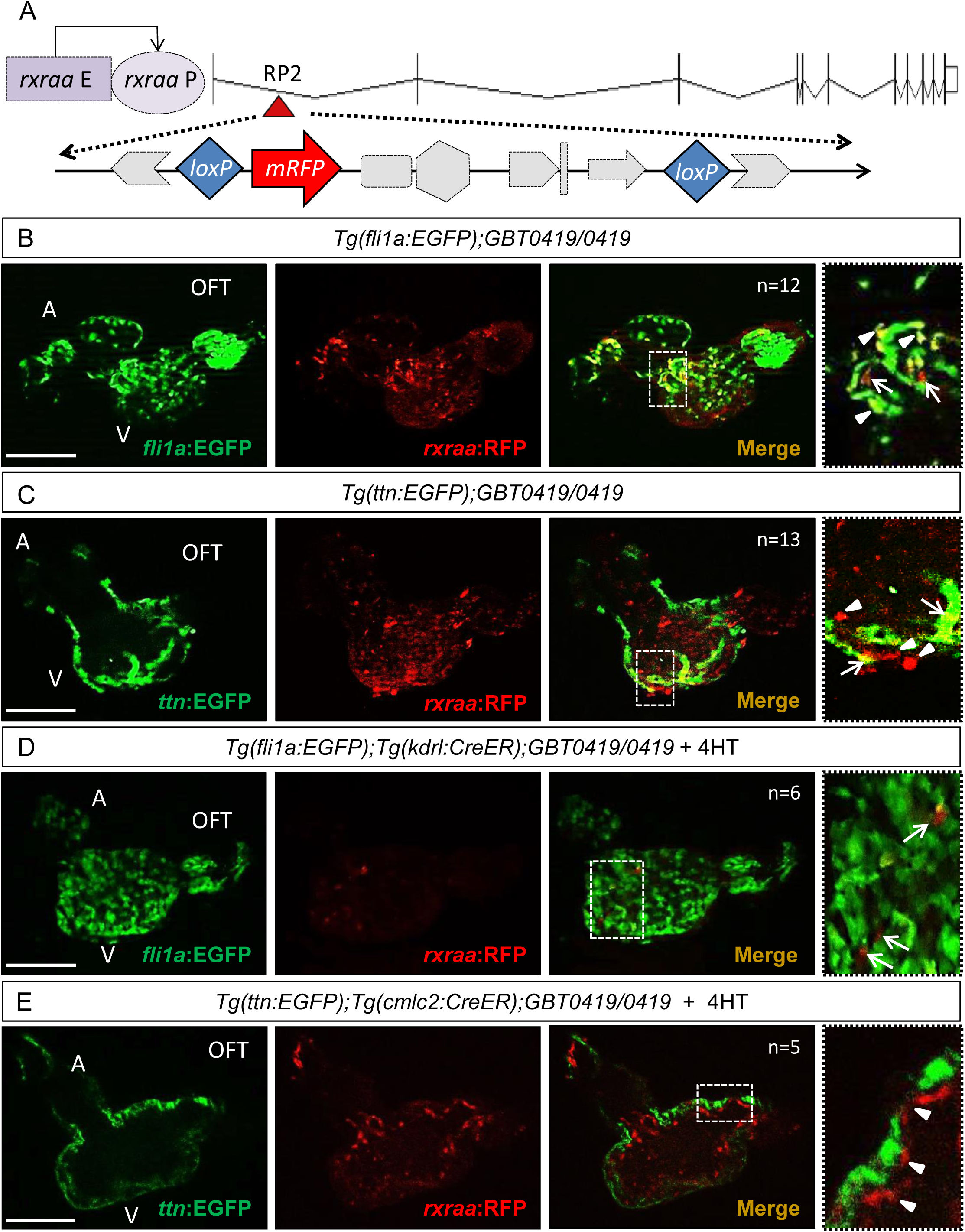
Endogenous *rxraa* expression in endothelial and myocardial cells of zebrafish hearts. **A**, Schematic of *rxraa* and location of the RP2 insertion (red triangle) in *GBT0419*. The RP2 vector contains an *mRFP* gene and 2 *loxP* elements. In *GBT0419*, RP2 is inserted in the first intron of *rxraa*. mRFP is fused in-frame with the N-terminal part of Rxraa and shows the expression driven by the endogenous *rxraa* enhancers. *rxraa* E, *rxraa* enhancers; *rxraa* P, *rxraa* promoter; mRFP, AUG-less monomeric red fluorescent protein. **B,** Images of a dissected heart from *Tg(fli1a:EGFP);GBT0419/0419* fish at 6 dpf. The far-right panel is a higher-magnification view of the dotted box area; triangles indicate mRFP expression in endothelial cells; arrows indicate non-endothelial expression of mRFP. **C,** Images of a dissected heart from *Tg(ttn:EGFP);GBT0419/0419* at 6 dpf. The far-right panel is a higher-magnification view of the dotted box area; arrows indicate mRFP expressed in cardiomyocytes; triangles indicate non-cardiomyocyte expression of mRFP. **D,** Image of a dissected heart of *Tg(fli1a:EGFP);Tg(kdrl:CreER);GBT0419/0419* fish at 6 dpf. Fish was treated by 4HT from 0 dpf to 6 dpf. The far-right is a higher-magnification view of the dotted box area; arrows indicate mRFP expression in non-endothelial cells. **E,** Image of a dissected heart of *Tg(ttn:EGFP);Tg(cmlc2:CreER);GBT0419/0419* fish at 6 dpf. Fish was treated by 4HT from 0 dpf to 6 dpf. The far-right panel is a higher-magnification view of the dotted box area with 90º rotation; triangles indicate non-cardiomyocyte expression of mRFP. In **(B)** to **(E)**, A, atrium; V, ventricle; OFT, out flow tract; 4HT, 4-hydroxytamoxifen. Scale bar=100 µm.

### Reversion of the RP2 insertion in endothelial cells abolishes the salutary effects of *GBT0419* allele on DIC

To further prove the salutary modifying effects of *GBT0419* are of cardiac nature and to discern whether endothelial or myocardial modulation of *rxraa* is vital for the modifying effects, we conducted cell-type specific rescue experiments by leveraging the integrated *loxP* sites in the RP2 vector (Figure 1A) (20). We bred *Tg(kdrl:CreER)* and *Tg(cmlc2:CreER)* into *GBT0419/0419* (homozygous mutant) to remove the RP2 insertion in the endothelium and myocardium, respectively (Figure 2A). We also generated a double-transgenic line with *Tg(tcf21:CreER)* to enquire potential functions of *rxraa* in the epicardium. To confirm the reversion of the RP2 insertion, we generated *Tg(fli1a:EGFP);Tg(kdrl:CreER);GBT0419/0419* and *Tg(ttn:actn-EGFP);Tg(cmlc2:CreER);GBT0419/0419* triple-transgenic lines. Indeed, after continuously treating embryos with 4-hydroxytamoxifen (4HT) from 0 to 6 dpf, the RP2 insertion could be effectively removed in a cell type-specific manner (Figure 1D, 1E), as indicated by the absence of mRFP expression in either endothelial cells or cardiomyocytes.

**Figure 2.**
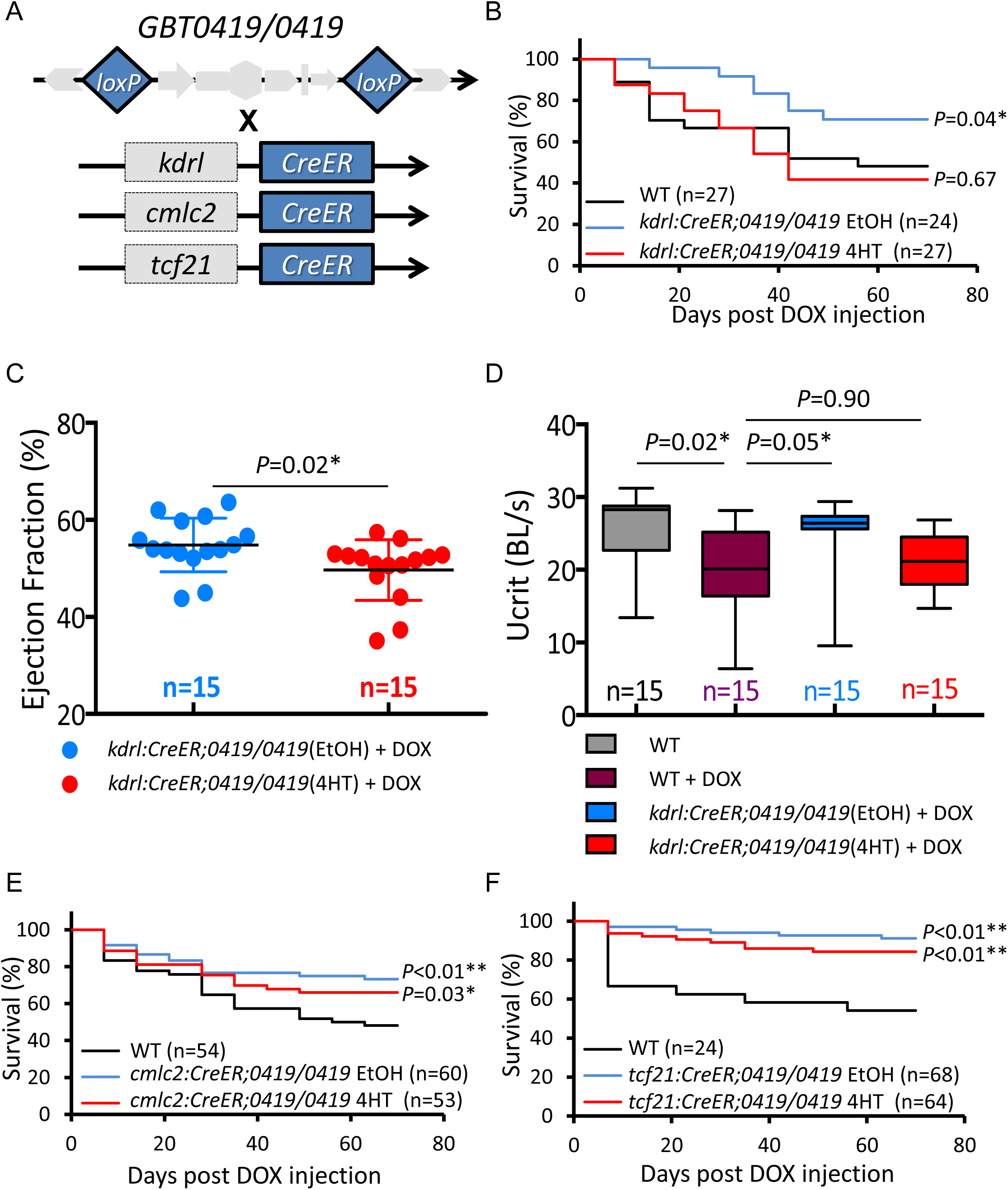
Endothelial RP2 reversion in *GBT0419/0419* abolishes its cardioprotective effects on DIC. **A,** Schematic shows the *Tg(kdrl:CreER), Tg(cmlc2:CreER)* and *Tg(tcf21:CreER)* transgenic lines that were crossed into *GBT0419/0419*, respectively. *CreER* and *loxP* elements were highlighted in blue. **B,** Kaplan-Meier survival curves indicate reduced survival of *GBT0419/0419* adult fish with endothelial RP2 reversion after DOX stress. **C,** Ventricular ejection fraction (EF) of *GBT0419/0419* adult fish was reduced with endothelial RP2 reversion after DOX stress. Each dot represent the EF value of a single ventricle. **D,** Swimming capacity of *GBT0419/0419* adult fish remains reduced with endothelial RP2 reversion after DOX stress. BL, body length; Ucrit, critical swimming speed. **E,** Kaplan-Meier survival curves indicate unchanged survival of *GBT0419/0419* adult fish with myocardial RP2 reversion after DOX stress. **F,** Kaplan-Meier survival curves indicate unchanged survival of *GBT0419/0419* adult fish with epicardial RP2 reversion after DOX stress. In **(B)**, **(E)** and **(F)**, WT, wild type; DOX, doxorubicin; EtOH, ethonal; 4HT, 4-hydroxytamoxifen. Error bars represent standard deviation. \**P*<0.05, \*\**P*<0.01; Log-rank test in **(B)**, **(E)** and **(F)**, shown statistics were comparisons to WT group; unpaired Student *t* test in **(C)**; 1-way ANOVA was used followed by posthoc tukey test in **(D)**.

We then treated 3 *Cre*-based double-transgenic embryos (Figure 2A) with either 4HT (to induce RP2 reversion) or ethanol (as a control), raised the fish up to 3 months old (Supplemental Figure 3A), stressed them with DOX, and assessed consequences of RP2 cell type-specific reversion in 3 ways. First, we monitored fish survival after DOX stress over a 10-week period. Consistent with our previous report (18), *GBT0419/0419* genetic background protects DOX-induced fish death (Figure 2B, 2E, 2F). Specific removal of the RP2 element in endothelial cells markedly diminished the salutary effects of *GBT0419/0419* on survival (Figure 2B). In contrast, RP2 reversion in cardiomyocytes and epicardial cells did not offset this survival-benefit (Figure 2E, 2F). Second, we assessed cardiac function of adult zebrafish using a recently developed *ex vivo* Langendorff-like system (44). While reduced ventricular ejection fraction was noted in wild type fish at 10 weeks post DOX injection (wpi), *GBT0419/0419* fish showed preserved pump function (Supplemental Figure 3B), indicating the salutary modifying effects on DIC is associated with a cardiac-specific protection. Similar to the survival index, reversion of the RP2 element in endothelial cells, but not in myocardial or epicardial cells, attenuated this cardioprotective effect (Figure 2C; Supplemental Figure 3D, 3E). Third, we quantified exercise capacity, a widely used clinical index for heart failure (45). We detected a gradual reduction in critical swimming speed (Ucrit) of adult fish with DIC at 4 wpi and thereafter (Supplemental Figure 3C), and noted a preserved Ucrit index in *GBT0419/0419* even at 10 wpi. This preserved swimming capacity was significantly attenuated by endothelial-specific removal of the RP2 insertion (Figure 2D). Collectively, these data indicated that the salutary modifying effects associated with *GBT0419/0419* on DIC are primarily attributable to the presence of the *GBT0419* allele in endothelial cells.

### The cardioprotective effects of the *GBT0419* allele is conferred by RA signaling activation

To gain insight into the molecular nature of the salutary effects of *GBT0419* allele, we generated *rxraa*^*e2*^, a transcription activator-like effector nucleases (TALEN) mutant containing an 8-nucleotide deletion in exon 2, the exon immediately following the RP2 inserted locus in *GBT0419* mutant (Figure 3A). This 8-nucleotide deletion will presumably lead to frameshift and result in truncated Rxraa proteins (Supplemental Figure 4). Surprisingly, *rxraa*^*e2/e2*^ failed to recapitulate salutary modifying effects of *GBT0419/0419* on DIC. The presence of *rxraa*^*e2*^ in the DIC model actually worsened the survival rate (Figure 3B). Whereas *GBT0419/0419* attenuated cardiomyocyte death, *rxraa*^*e2/e2*^ showed a higher ventricular cardiomyocyte apoptosis index at 8 wpi (Figure 3C, 3D). In contrast to rescued cardiac function in *GBT0419/0419* mutant, we observed no difference between *rxraa*^*e2/e2*^ mutant and wild type control after DOX stress, with both groups showed increased ventricular end-systolic volume (ESV) and reduced ventricular ejection fraction (EF) (Figure 3E, Supplemental Figure 5). We measured the expression of 2 pathological markers of cardiac remodeling, *natriuretic peptide A* (*nppa)* and *natriuretic peptide B* (*nppb*) (Figure 3F, 3G), and detected induction of *nppa* in *rxraa*^*e2/e2*^ mutant but not in *GBT0419/0419*. Whereas myofibril organization in *GBT0419/0419* at 12 wpi was characterized by more even distribution of α-actinin expression, improved lateral alignment and better-organized sarcomeres (compared to wild type control group), in *rxraa*^*e2/e2*^, myofibril remained disorganized (Figure 3H). Collectively, these data strongly indicated *GBT0419/0419* is exerting a different mechanism than a simple loss-of function outcome at the *rxraa* locus.

**Figure 3.**
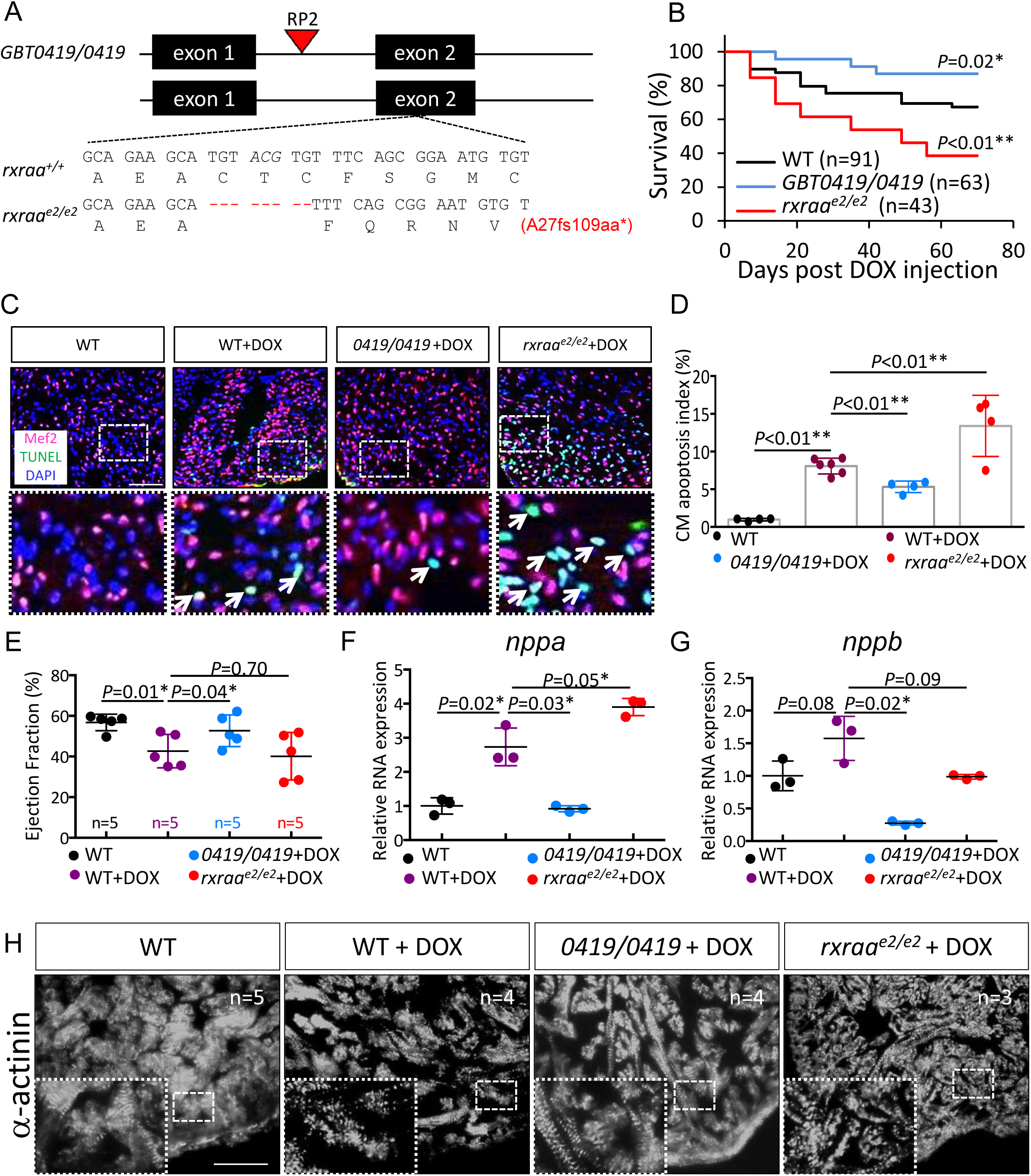
While *GBT0419* exerts salutary effects, a *rxraa* TALEN allele exerts deleterious effects on DIC in adult zebrafish. **A,** Schematic comparison of genomic lesions in *GBT0419/0419* and a *rxraa* TALEN mutant (*rxraa*^e2/e2^). Dashes indicate an 8 nucleotides deletion. fs, frameshift; aa, amino acid. **B,** Kaplan-Meier survival curves show survival of WT, *GBT0419/0419* and *rxraa*^e2/e2^ adult fish after DOX stress. **C,** Representative images of cardiomyocyte apoptosis in ventricular tissues of WT, *GBT0419/0419* and *rxraa*^e2/e2^ after DOX stress at 10 wpi. The lower panel is higher-magnification view of the corresponding dotted box area. Arrows indicate cardiomyoctye (Mef2^+^) with positive TUNEL staining. Mef2, myocyte enhancer factor 2; TUNEL, terminal deoxynucleotidyl transferase dUTP nick end labeling; DAPI, 4’,6-diamidino-2-phenylindole. Scale bar=50 µm **D,** Quantification of cardiomyocyte apoptosis index of **(C)**. Each dot represents a single fish ventricle. About 500-1,000 cardiomyocytes were counted for sections per ventricle. CM, cardiomyocyte. **E,** Ventricular ejection fraction of WT, *GBT0419/0419* and *rxraa*^e2/e2^ after DOX stress at 10 wpi. **F, G,** Shown are expression of *nppa* **(F)** and *nppb* transcripts **(G)** in WT, *GBT0419/0419* and *rxraa*^e2/e2^ after DOX stress at 10 wpi. RNA was extracted from a pool of 3 ventricles as a single biological replicate. n=3 in each group. **H,** Images of *α*-actinin antibody staining shows myofibril disarray in ventricular tissues of WT, *GBT0419/0419* and *rxraa*^e2/e2^ after DOX stress at 10 wpi. Insets are higher-magnification views of dotted box areas. Scale bar=50 µm. WT, wild type; DOX, doxorubicin. Error bars represent standard deviation. \**P*<0.05, \*\**P*<0.01; Log-rank test in **(B),** shown statistics were comparisons to WT group; 1-way ANOVA followed by posthoc Tukey test in **(D)**, **(E)**, **(F)** and **(G)**.

We then decided to carefully look into molecular altering between *GBT0419* and *rxraa*^*e2*^ mutants. In *GBT0419/0419,* splicing between exon 1 and 2 of *rxraa* was disrupted by >90%, as revealed by the primer pair F1-R1 for quantitative PCR, indicating high protein-trap efficiency (Figure 4A, 4B). The expression levels of chimeric *rxraa* transcripts containing exons 2, 3 and 4, as revealed by the primer pair F2-R2, are 3 times higher than those in wild type siblings (Figure 4A, 4B, 4C). This is expected since the RP2 vector includes strong enhancer/promoter driving stabilized GFP expression upon successful 3’ splicing of a chimeric transcript (20). Most chimeric transcript shall not encode any functional endogenous proteins, because of a stop codon after the GFP encoding sequence in RP2 vector. However, we noted an in-frame alternative start codon in exon 2 of *rxraa* that is located after the RP2 insertion (Supplemental Figure 1A), raising the possibility of a gain-of function mechanism for *GBT0419* mode of action. Different from *GBT0419/0419*, expression level of *rxraa* transcripts was reduced by ∼50% in *rxraa*^*e2/e2*^ (Figure 4B, 4C). The levels of *rxrab* transcripts remained unchanged in both mutants (Supplemental Figure 6A).

**Figure 4.**
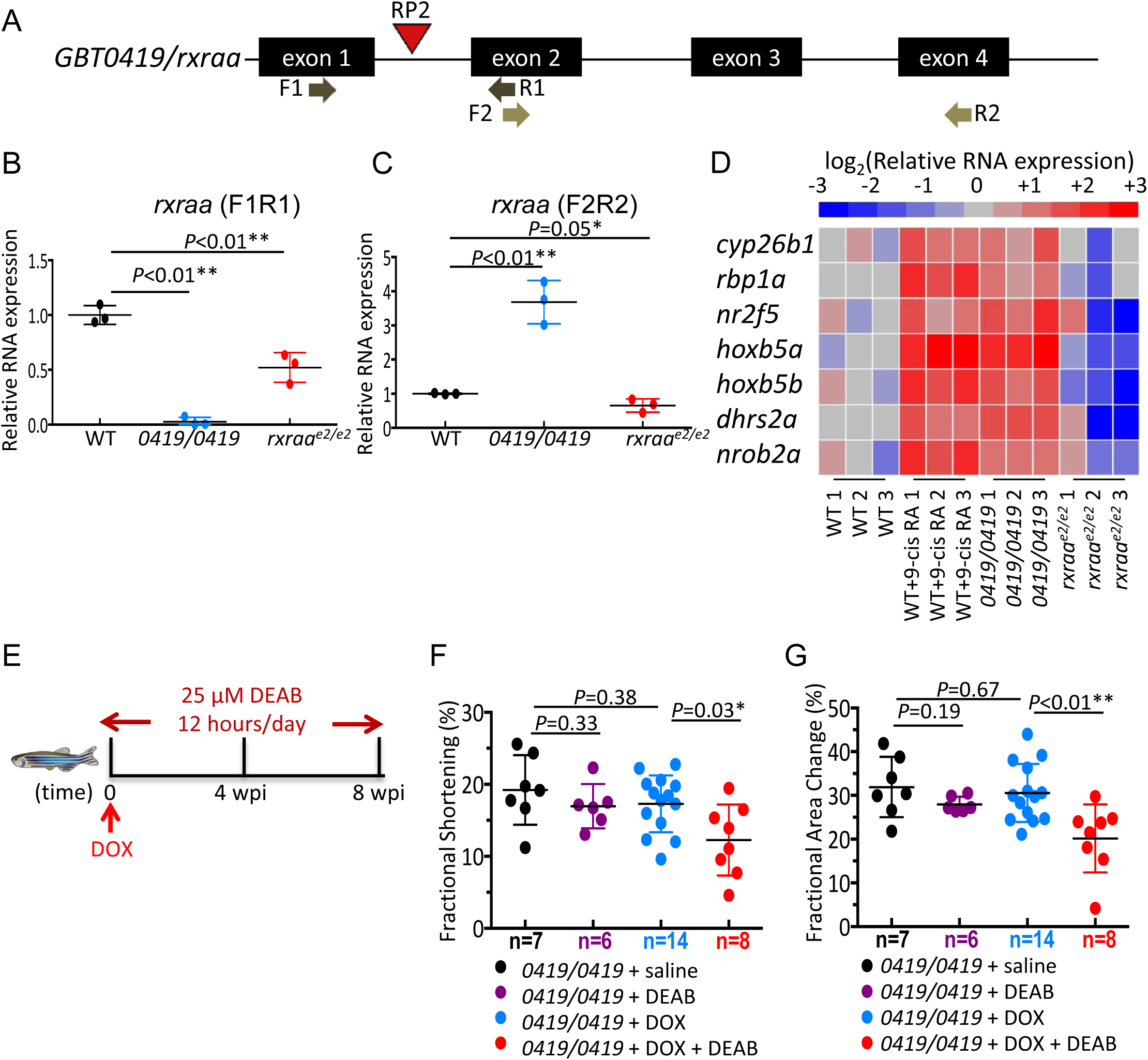
Activation of RA signaling is responsible for the cardioprotective effects associated with *GBT0419*. **A,** Schematics showing locations of *rxraa* primers for quantitative RT-PCR. Primer pair 1 (F1R1) targets on exon 1 and exon 2, flanking the RP2 insertion locus. Primer pair 2 (F2R2) targets on exon 2 and exon 4, which shall amplify *rxraa* transcripts containing C-terminal exons. **B,** Relative degree of exon 1 - exon 2 splicing *rxraa*, as measured by primer pair 1, in *GBT0419/0419* and *rxraa*^e2/e2^. **C,** Relative expression of *rxraa* transcripts, as measured by primer pair 2, in *GBT0419/0419* and *rxraa*^e2/e2^. **D,** Heatmap of relative mRNA expressions levels of genes regulated by RA signaling in *GBT0419/0419* and *rxraa*^e2/e2^. Values are shown as log2 transformation of relative mRNA expression normalized to the mean value of WT. *gapdh* was used as an internal control. In **(B)** to **(D)**, RNA were extracted from a pool of 20 fish embryo at 2 dpf and considered as a single biological replicate (represented as a single dot). Samples were collected in triplicates. **E,** Schematic of the schedule for DEAB administration and DOX injection. wpi, weeks post DOX injection. **F,** Ventricular fractional shortening of *GBT0419/0419* adult fish, with or without DEAB treatment, upon DOX stress at 8 wpi. **G,** Ventricular fractional area change of *GBT0419/0419* adult fish, with or without DEAB treatment, upon DOX stress at 8 wpi. Each dot represents an individual ventricle. WT, wild type; 9-cis RA, 9-cis retinoic acid; DEAB, diethylaminobenzaldehyde. Error bars indicate standard deviation. \**P*<0.05, \*\**P*<0.01; 1-way ANOVA followed by posthoc Tukey test was used.

To test the gain-of function hypothesis, we explored *rxraa* expression at protein and activity levels. We tried several antibodies against mouse Rxra in zebrafish; unfortunately, none were sufficiently specific for either Western blotting or immunostaining analysis. As an alternative approach, we assessed RA signaling activity by measuring transcriptional expression of several known RA-targeted genes (46-48). As a positive reference state for these studies, exogenous treatment with 9-cis RA on 2 dpf wild type zebrafish embryos resulted in an increase in the transcription of these RA-targeted genes. By comparison, most of these gene transcripts were significantly induced in *GBT0419/0419*, but inhibited or unchanged in *rxraa*^*e2/e2*^ (Figure 4D, Supplemental Figure 6D). These data supported the gain-of function hypothesis that, different from the *rxraa*^*e2/e2*^indel mutant, RA signaling is net activated in *GBT0419/0419*.

To further test whether such activated RA signaling is the molecular basis underlying the salutary modifying effects of *GBT0419/0419*, we inhibited RA signaling with diethylaminobenzaldehyde (DEAB) (46, 49, 50) (Figure 4E), a compound able to inhibit the activity of Aldehyde dehydrogenase 1 family, member A2 (Aldh1a2), the major enzyme for endogenous RA synthesis. We found treatment of DEAB worsened the survival of *GBT0419/0419* upon DOX stress (Supplemental Figure 7A), and attenuated its cardioprotective effects at both 4 wpi (Supplemental Figure 7B, 7C, 7D) and 8 wpi (Figure 4F, 4G), as evidenced by reduced ventricular pump function measured by high-frequency echocardiography (51). Together, these data supported the conclusion that the activation of RA signaling through a gain-of-function mechanism in the *GBT0419* allele confers its salutary modifying effects on DIC.

### Endothelial-specific overexpression of *rxraa* is therapeutic to DIC

To directly determine whether RA signaling activation, through gain-of function of *rxraa*, is therapeutic for DIC, we generated an inducible transgenic line for the gene, *Tg(βactin2:loxP-mCherry-stop-loxP-rxraa-EGFP)*, hereafter termed *Tg(βact2:RSRxraa)* (Figure 5A). To test the binary expression system, we crossed *Tg(βact2:RSRxraa)* into *Tg(hsp70l:Cre)* and stressed the double-transgenic fish by heat shock at the embryonic stage. We noted EGFP fluorescence (Supplemental Figure 8A), confirmed the identity of the Rxraa-EGFP protein by Western blotting with an antibody against EGFP (Supplemental Figure 8B), and detected activation of RA-targeted genes (Supplemental Figure 8C), indicating successful overexpression of a functional chimeric protein that activates RA signaling pathway.

**Figure 5.**
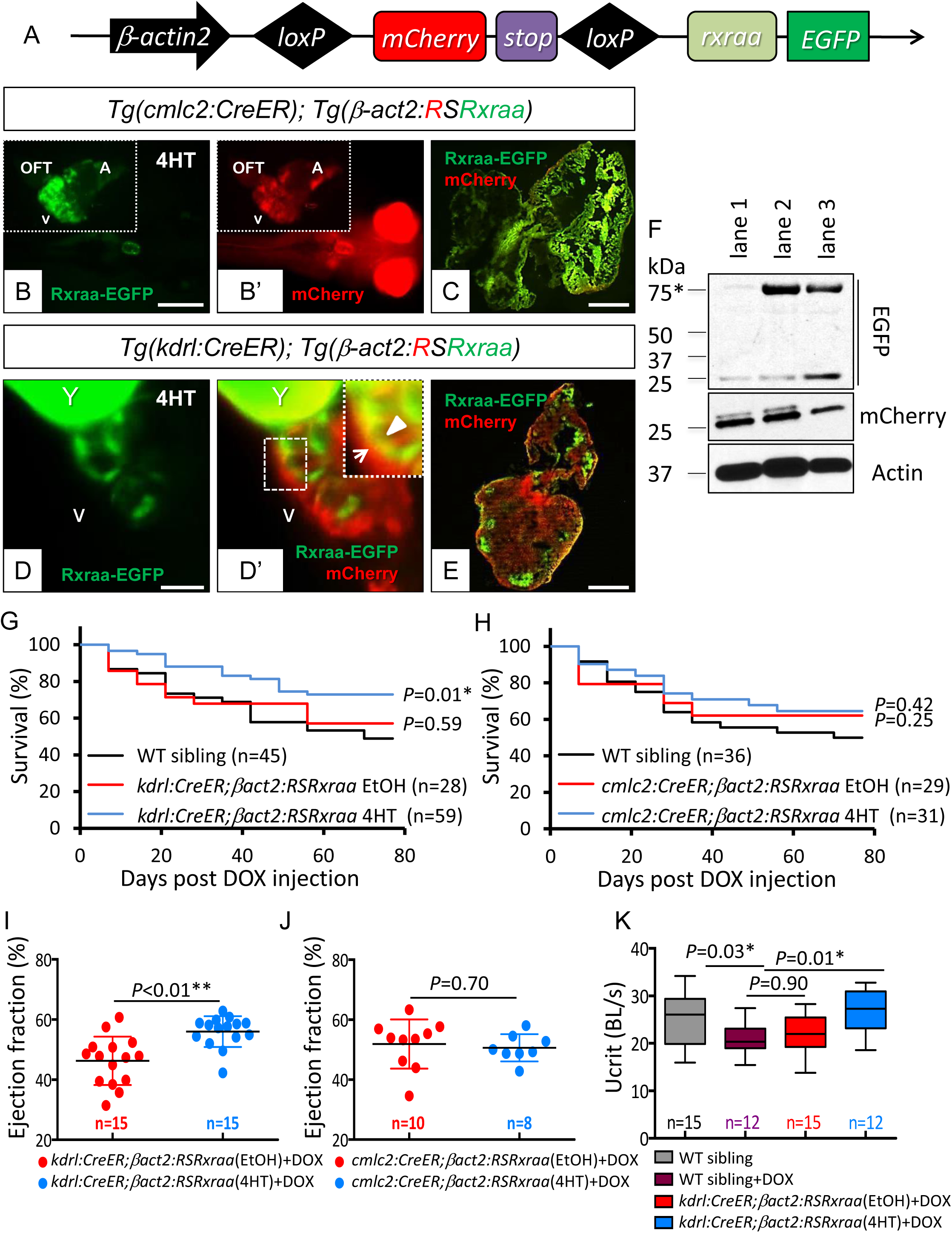
Endothelial-specific overexpression of *rxraa* protects against DIC *in vivo*. **A,** Schematic of the *rxraa* conditional transgenic line. **B,** The green channel shows cardiomyocyte-specific *rxraa* overexpression at 6 dpf. **B’,** The red channel of the same view as **(B)** shows the original mCherry expression. Insets show the same heart after dissection. **C,** Heart from a 3-month-old fish showing cardiomyocyte-specific *rxraa* overexpression. Scale bar=500 µm. **D,** The green channel shows endothelial-specific *rxraa* overexpression at 6 dpf. **D’** The merged channel with the same view as **(D)** shows mCherry and Rxraa-EGFP expressions. Inset shows a higher-magnification view of the dotted area. Triangle indicates the endothelial layer; arrow indicates the non-endothelial layer. Scale bar=100 µm. **E,** Heart from a 3-month-old fish showing endothelial-specific *rxraa* overexpression. Scale bar=500 µm. In **(B)** to **(E)**, A, atrium; V, ventricle; OFT, out flow tract; Y, yolk with auto-florescence. **F,** Western blot of adult ventricles. Predicted size of the induced Rxraa-EGFP fusion protein is 75 KDa (indicated by an asterisk). Actin is used as an internal control. Lane 1, *Tg(βact2:RSRxraa)*; lane 2, *Tg(βact2:RSRxraa);Tg(kdrl:CreER)* + 4HT; lane 3, *Tg(βact2:RSRxraa);Tg(cmlc2:CreER)* + 4HT. **G, H,** Kaplan-Meier survival curves showing survival of DOX-stressed adult fish with endothelial-specific **(G)** or cardiomyocyte-specific **(H)** *rxraa* overexpression. **I, J,** Ventricular ejection fraction of adult hearts with endothelial-specific **(I)** or cardiomyocyte-specific **(J)** *rxraa* overexpression at 8 wpi. **K,** Swimming capacity of fish, with or without endothelial-specific *rxraa* overexpression. Ucrit, critical swimming speed; BL, body length. WT, wild type; DOX, doxorubicin; EtOH, ethonal; 4HT, 4-hydroxytamoxifen; wpi, weeks post DOX injection. Error bars indicate standard deviation. \**P*<0.05, \*\**P*<0.01; Log-rank test in **(G)** and **(H)**; unpaired Student *t* test in **(I)** and **(J)**; 1-way ANOVA followed by posthoc Tukey test in **(K)**.

We then bred *Tg(βact2:RSRxraa)* into *Tg(cmlc2:CreER)* and *Tg(kdrl:CreER)* to assess the effects of Rxraa overexpression in myocardial and endothelial cells, respectively. Effective induction of chimeric protein in either cardiomyocytes or endothelial cells after 4HT treatment was noted from embryogenesis (Figure 5B, 5D) through adulthood (Figure 5C, 5E). The identity of the Rxraa-EGFP fusion protein in adult cardiac tissues was further confirmed by Western blotting (Figure 5F). We then stressed these 4HT-induced double-transgenic fish with DOX and showed that endothelial-specific overexpression of *rxraa* was therapeutic to DIC, as indicated by rescued ventricular function, restored exercise capacity (Figure 5I, 5K), and improved survival (Figure 5G). In contrast, cardiomyocyte-specific *rxraa* overexpression did not exert any beneficial effects on DIC (Figure 5H, 5J).

We also evaluated effects of epicardial *rxraa* overexpression on DIC by generating *Tg(tcf21:CreER); Tg(βact2:RSRxraa)* double-transgenic fish. Upon 4HT treatment, we could detect some Rxraa-EGFP fusion protein induction in embryonic *tcf21+* cells (6 dpf) at the junction between the outflow tract (OFT) and the ventricle (Supplemental Figure 9A). We did not note any protective effects on DIC from this double-transgenic line (Supplemental Figure 9C). Of note, these data should be cautiously interpreted because a previous report (52) suggested that the *βactin2* promoter might not drive gene expression efficiently in the *tcf21*^*+*^ epicardial cells. We also did not detect consistent induction in adult fish hearts at 3 months old (Supplemental Figure 9B).

Together with the conditional rescue experiments, our transgenic analysis provided strong *in vivo* evidence supporting the conclusion that overexpression of *rxraa* results in RA signaling activation and exerts therapeutic benefits on DIC. Importantly, endothelial-specific overexpression of *rxraa* is exclusively vital for such therapeutic benefits.

### RXRA agonists exert stage-dependent therapeutic effects on DIC

Having confirmed RA signaling activation as the underlying mechanism of salutary *rxraa* alleles, we then sought to investigate if RXRA agonists could be potentially used to treat DIC. We first recapitulated a high throughput embryonic zebrafish DIC model (53), i.e. administration of DOX in wild type embryos from 24 to 72 hours post fertilization (hpf). DIC phenotypes, including reduced ventricular shortening fraction (SF), heart rate and survival, could be effectively rescued in *GBT0419/0419* (Supplemental Figure 10), indicating the feasibility of applying this model to assess RA-based therapies. We then assessed therapeutic potentials of 6 commercially available compounds (Table 1) that activate RXRA (54-59). To exclude teratogenic effects associated with RA signaling activation (60-62), we experimentally determined the median lethal dose (LD50) of each compound (Table 1, Supplemental Figure 11A), and then titrated doses. Teratogenic effects, such as pericardial edema (Supplemental figure 11B), can be noted for 4 compounds when treated at doses ranging from 20%-50% LD50, but not for any of these 6 RXRA agonists when administered around 1% LD50 (Table 1). At these safe doses, we observed therapeutic effects of isotretinoin, SR11237 and bexarotene on DIC, as indicated by reduced mortality (Figure 6C), improved ventricular shortening fraction and restored heart rate (Figure 6D, 6E).

**Table 1.** Dose determination for 6 RXRA agonists in zebrafish embryos.

| **RXRA agonist <sup>1</sup>** | **LD50 (95% CI)**<br>**μM** | **Cardio-phenotypic dose <sup>2</sup>**<br>**μM** | **Doses used in Figure 6 <sup>3</sup>**<br>**μM** |
| --- | --- | --- | --- |
| ATRA | 2.82 (2.26-3.42) | 0.2 | 0.028 |
| 9-cis RA | 3.14 (2.71-3.55) | 1.0 | 0.031 |
| DAM | 0.99 (0.86-1.15) | N/A | 0.010 |
| isotretinoin | 3.84 (2.79-5.23) | 0.6 | 0.038 |
| SR11237 | 4.69 (3.05-5.16) | N/A | 0.047 |
| bexarotene | 4.26 (0.31-5.27) | 4.0 | 0.043 |

<sup>1</sup> RXRA agonists were added to water with embryos from 24 hpf to 72 hpf; phenotypes were analyzed at 72 hpf. Embryonic lethal was defined as the cessation of heart contractions.
<sup>2</sup> Cardio-phenotypic dose was defined as the minimal dose that caused any deleterious cardiac phenotypes (**Supplemental Figure 10B**).
<sup>3</sup> Compounds at the dose of 1% LD50 were used in **Figure 6**.
Abbreviations: CI, confidence interval; hpf, hours post fertilization; ATRA, all trans retinoic acid; 9-cis RA, 9-cis retinoic acid; DAM, docosahexaenoic acid. LD50, median lethal dose; NA, not applicable.

**Figure 6.**
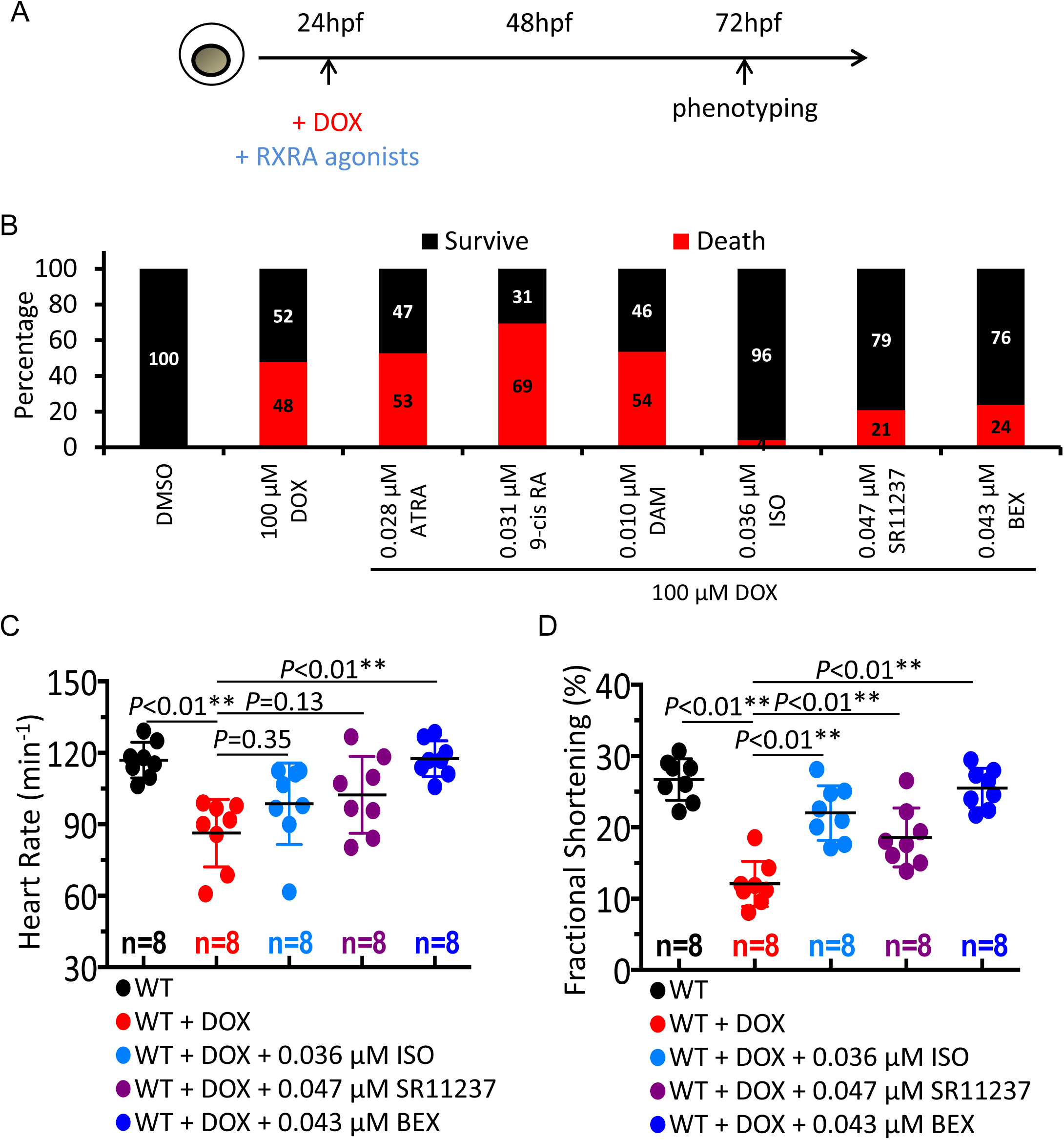
Assessment of cardioprotective effects of 6 RXRA agonists using an embryonic zebrafish DIC model. **A,** Schematic of the schedule for assessing effects of RXRA agonist treatment. hpf, hours post fertilization. **B,** Survival of embryos at 72 hpf. Embryos were cotreated with 100 µM DOX and various RXRA agonists at indicated doses. At least 36 embryos were used in each group. **C,** Heart rate in embryos stressed with 100 µM DOX alone or plus ISO, SR11237 or BEX. **D,** Fractional shortening in embryos stressed with 100 µM DOX alone or plus ISO, SR11237 or BEX. In (**B**) to (**D**), WT, wild type; DOX, doxorubicin; ATRA, all trans retinoic acid; 9-cis RA, 9-cis retinoic acid; DAM, docosahexaenoic acid; ISO, isotretinoin; BEX, bexarotene. Error bars indicate standard deviation. \**P*<0.05, \*\**P*<0.01; 1-way ANOVA followed by posthoc Tukey test was used in **(C)** and **(D)**.

Next, we selectively assessed therapeutic effects of isotretinoin and bexarotene, 2 FDA approved drugs, using our adult zebrafish DIC model. Given this model progresses from an early acute phase with normal cardiac function to a late chronic phase with reduced cardiac dysfunctions at around 4 weeks post injection (wpi) (16), we assessed efficacy of these 2 RXRA agonists in 3 different phases, i.e. 1 week prior to DOX administration (pretreatment), 1-4 weeks post DOX injection (early stage DIC), and 5-8 weeks post DOX injection (late stage DIC) (Figure 7A, 7E, Supplemental Figure 12D). Both compounds were administered to adult fish via daily oral gavage (63, 64). The doses (24 µg/day/fish for isotretinoin or 97 µg/day/fish for bexarotene) were inferred from the FDA-recommended oral doses for human cancer patients (Supplemental Table 2). At these doses, a 1-time gavage of either chemical was able to activate *rxraa* in adult fish hearts within 24 hour (Supplemental Figure 12A). Interestingly, we found such treatment of either RXRA agonist (on daily basis) was therapeutically effective only when administered during early stage DIC, as shown by increased survival (Figure 7B) and improved cardiac function at 4 wpi (Supplemental Figure 12B, 12C). Although we stopped treatment at 5 wpi and thereafter, survival remained enhanced from 5 wpi to 8 wpi, with preserved heart function (Figure 7C, 7D). In contrast, pretreatment for 1 week failed to improve cardiac function at 4 wpi (Supplemental figure 12E, 12F), and treatment at the late stage DIC could even have detrimental effects, as indicated by the worse survival rate (compared to wild type controls) with marginal statistical significance (Figure 7F, *P*=0.16 and *P*=0.13 respectively, Log-rank test) and unchanged cardiac functions at 8 wpi (Figure 7G, 7H). Of note, we detected mild induction of *rxraa* and *rxrab* transcripts in the ventricular tissue at the early phase post DOX stress and much stronger induction of these 2 transcripts at the later phase (Supplemental Figure 12G). Together, we concluded that RA signaling activation might represent a compensational pathway along DIC progression, which could be modulated (by treatments of RXRA agonists) for therapeutic benefits in a temporally restricted manner.

**Figure 7.**
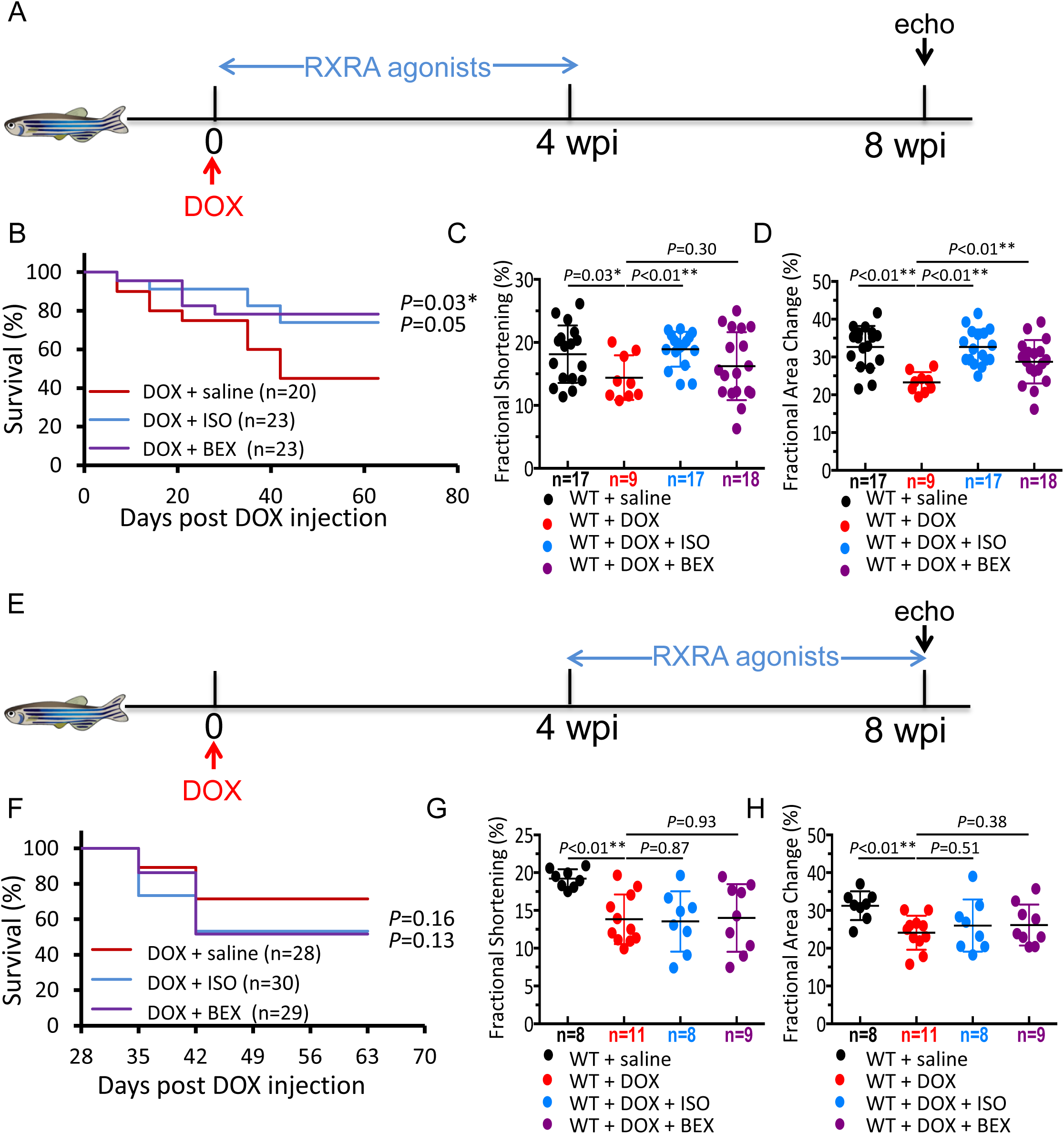
RXRA agonists exert stage-dependent cardioprotective effects on DIC. **A, E,** Schematics of the schedules for RXRA agonist treatment at early (1 wpi to 4 wpi) or late (4 wpi to 8 wpi) stages of the adult fish DIC model. A single bolus of DOX was injected as indicated. RXRA agonists were delivered daily by oral gavage. High frequency echocardiography was performed at the indicated time points to quantify cardiac functions. Saline was used as a control. wpi, weeks post DOX injection. **(B)**, **(C)** and **(D)** show an early phase treatment with RXRA agonists. **(F)**, **(G)** and **(H)** show late phase treatment with RXRA agonists. **B, F,** Kaplan-Meier survival curves show survival of adult fish after DOX stress. **C, G,** Fractional shortening of adult ventricles at 8 weeks post DOX stress. **D, H,** Fractional area change of adult ventricle at 8 weeks post DOX stress. ISO and BEX were administered at clinically relevant doses as listed in the **Supplemental Table 2**. WT, wild type; DOX, doxorubicin; ISO, isotretinoin; BEX, bexarotene. Error bars represent standard deviations. \**P*<0.05, \*\**P*<0.01; Log-rank test was used in **(B)** and **(F),** shown statistics were comparisons to saline group; 1-way ANOVA followed by posthoc Tukey test was used In **(C)**, **(D)**, **(G)** and **(H)**.

To further explore the endothelial nature of the cardioprotective effects of isotretinoin and bexarotene, we tested both compounds on primary human coronary artery endothelial cells *in vitro*. We showed that isotretinoin effectively prevented DOX-induced endothelial cell death (Supplemental Figure 13A), and treatment of either RXRA agonist stimulated endothelial paracrine signaling (Supplemental Figure 13B, 13C) that have been previously recognized with therapeutic benefits on heart failure (65-68). Notably, endothelial nitric oxide synthase (eNOS) signaling was found to be a common pathway that was activated by treatments of both compounds. These data suggested that activating RA signaling in endothelium could promote cell viability and enhance secretion of salutary molecules, both of which might contribute to the associated cardioprotective effects.

## Discussion

### *rxraa* is a therapeutic gene target for DIC in adult zebrafish

Here we conducted comprehensive series of studies on *GBT0419,* the first salutary modifying mutant emerged from a mutagenesis screen, and reported *rxraa* as a gene target for cardioprotection against DIC. We proved the cardiac nature of the modifying effects, and further demonstrated that *GBT0419*-associated cardioprotective effects was conferred, at least in part, by the activation of RA signaling, which could be achieved by either *rxraa* overexpression or administration of RA activators. We subsequently identified 2 FDA-approved RXRA agonists, bexarotene and isotretinoin, that could be used to treat zebrafish DIC. These candidate compounds could be considered in future preclinical studies, with the goal of repurposing them for treating DIC patients.

The aforementioned “gain-of function” outcome in *GBT0419* appears to be different from what have be previously described in other GBT lines, as all have been to date demonstrably loss-of-function mutants (17, 20, 40, 41, 69). However, it is still possible that *GBT0419* is a loss-of function allele at mRNA or even protein level, given that this mutant has a blunted *rxraa* response to 9-cis RA (Supplemental Figure 6B, 6C), and the activated RA signaling in *GBT0419* could be the consequence of certain ‘feed-forward’ mechanism that has been previously recognized (47). Nevertheless, the strong genetic and pathway altering data are all consistent with *GBT0419* is an *rxraa* gain-of function allele at the RA signaling level.

Instead of further digging the intricate molecular nature associated with the GBT-based protein-trap, we focused on experiments that directly sought to characterize therapeutic insights underlying *RXRA* manipulation for DIC in current study. First, we showed that DEAB, an inhibitor for RA synthesis, largely depletes the cardioprotective effects of *GBT0419/0419*. Second, we generated tissue-specific transgenic fish lines and demonstrated cardioprotective effects of *rxraa* endothelial-specific overexpression. Unlike mice with cardiomyocyte-specific overexpressed *Rxra* (35, 36), the transgenic fish with cardiomyocyte-specific overexpression of *rxraa* did not have any noticeable cardiac phenotype. It is possible that the much stronger regenerative capacity of the zebrafish heart have rendered it more resilience to the biomechanical stress. Third, we showed that daily administration of 2 RXRA agonists could exert cardioprotective effects against DIC in adult zebrafish. Together with those previous reported pharmacological evidence (23-30), our study provided the first *in vivo* genetic evidence in support of *RXRA* as a therapeutic target for DIC, which functions by activating RA signaling.

### An endothelial cell- and early phase-specific guideline could be considered in future translations of RXRA-based therapies

The tissue-specific analysis of the DIC modifying effects of *rxraa* in adult zebrafish revealed a novel therapeutic opportunity by targeting *RXRA* in the endothelium. It is plausible that endothelial *RXRA* was not recognized as a therapeutic target because previous functional studies of *RXRA* were primarily conducted in cardiomyocytes (35, 36). If gene-based therapy is pursued, an endothelial-specific promoter could be integrated in its molecular designs. When compound-based therapy is pursued, technologies that facilitate endocardium-preferred drug delivery should be considered (70-72). Given endothelial dysfunction is a known pathological characteristic of various cardiovascular diseases including DIC (65, 73-75), deeper mechanistic studies of RXRA-based therapies are worthy to be pursued. Our cell culture data supported that treatment of RXRA agonists could directly counteract doxorubicin-induced toxicity in endothelial cells (Supplemental Figure 13A), and/or indirectly regulating signaling pathways in cardiomyocytes via modifying secreted molecules such as nitric oxide, Neuregulin 1 and Fibroblast growth factors along the lines of endothelial-myocardial crosstalk (66-68)(Supplemental Figure 13B, 13C). Additionally, potential contribution from the regenerative mechanisms guided by RA signaling shall be assessed (43).

In addition to the spatial specificity of the RXRA-based therapies, we also uncovered its temporal specificity, i.e. RXRA agonists are therapeutic only if administered in the early phase of DIC, but not in the later phase when cardiac dysfunction becomes apparent or in the phase before DOX exposure. By tracking gene expression in the adult zebrafish hearts, we observed transcriptional upregulation of *rxraa* and *rxrab* began at 1 week after DOX stress, and was considerably more activated by 8 weeks (Supplemental Figure 12G). Together with the early therapeutic time window, we postulate that RA signaling activation represents a compensational response to DOX stress in the heart. Given that reduction of *RXRA* expression has been previously detected in heart failure cohorts and a canine model of heart failure (76, 77), as well as in a rat model of diabetic cardiomyopathy (29, 30), future studies are warranted to determine whether RXRA-based therapies can also be leveraged in different types of cardiovascular diseases. Notably, a temporal-specific effect has also been elucidated for mammalian target of rapamycin (mTOR)-based therapies in our previous studies using the same model (16). In contrast to RA signaling activation, mTOR inhibition exerts therapeutic effects only in the later phase of DIC. Future comparisons of RA- and mTOR-based therapies should uncover additional insights on stage-dependent mechanisms of DIC, and might lead to combinatory therapeutic strategies.

Because of the pleiotropic functions of RA signaling, the impact of RA-based therapies on other organs needs to be considered carefully. Of note, RA-receptor(s) agonists are being actively developed as anticancer agents (78), and combination chemotherapies with DOX have been explored in clinical trials (79). Collaborations among scientists with expertise in different area will ultimately accelerate translation of RXRA-based therapies from bench to bedside in a safe and prudent manner.

### Forward genetic screening in zebrafish can be leveraged to rapidly identify novel therapeutic targets for cardiomyopathies

The characterization of *GBT0419*, the first salutary mutant identified from our forward genetic screen, established the feasibility of taking phenotype-based screening approaches to effectively discover therapeutic targets for cardiomyopathies. We believe this innovative application can be highly significant because traditional approaches might easily overlook the therapeutic capacity of a candidate gene if it is not manipulated precisely as needed. Meanwhile, our data demonstrated the efficiency of zebrafish models for follow-up mechanistic studies. The integrated *loxP* sites in all GBT mutants and ability to simultaneously analyze multiple double- or triple-transgenic fish lines facilitate the elucidations of any spatially specific therapeutic effects. The relatively high throughput of zebrafish models also accelerates comparison of multiple drugs, identification of most effective candidates, and optimization of treatment timing.

Our work justifies future scaling of the zebrafish-based modifier screen to the whole genome for systematic discovery of more genetic modifiers. Inferred from our pilot screen, it is estimated that ∼200 genetic modifiers for DIC exist (18), and identification of additional myocardial-, endothelial-, or potentially epicardial-based modifiers will expedite a comprehensive understanding of the pathogenic processes that lead to cardiomyopathy. Salutary modifiers will be studied with higher priority, as each of which might directly lead to a new therapeutic avenue for human patients at risk of or with developing DIC. It is also highly expected that some of these therapeutic strategies can also be extended to other types of cardiomyopathies and/or heart failure.

## Methods

### Zebrafish husbandry

Adult zebrafish were maintained under a light-dark cycle (14 hours of light, 10 hours of darkness) at 28.5 °C. Zebrafish embryos were maintained in 10-mm petri dish with E3 water at 28.5 °C up to 7 dpf. All animal experiments were approved by the Mayo Clinic College of Medicine Institutional Animal Care and Use Committee (protocol A00002783-17).

### Generation of the *rxraa*^*e2/e2*^ mutant

Transcription activator-like effector nucleases (TALENs) targeting exon 2 of *rxraa* were designed with Zifit (http://zifit.partners.org/ZiFiT/ChoiceMenu.aspx). The N-terminus TALEN-binding sequence was 5’-tcagagaggacgagcaga-3’, and the C-terminus TALEN-binding sequence was 5’-cggaatgtgtgctcttca-3’. Both TALENs were constructed by using the Golden Gate kit (80). Capped mRNAs were synthesized by using mMESSAGE mMachine T3 kit (Ambion). TALEN mRNAs (15 pg to 1 ng) were injected into wild type embryos at 1-cell stage. Founder fish with desired genomic lesion were identified by genotyping (forward primer, 5’-tccatagggttcctgaagc-3’; reverse primer, 5’-tctgaatatgcacgttacctg-3’). The amplified PCR products were digested with restriction enzyme HypCH4IV. The precise genomic lesion was then determined by Sanger sequencing. After the mutants were generated and sequenced, the founders were outcrossed to WIK fish for >5 generations to eliminate any potential background mutations.

### Generation of the *Tg(βactin2: loxP-mCherry-stop-loxP-rxraa-EGFP)* transgenic line

The transgenic line was generated using Tol2-gataway system (81). The *loxP-mCherry-stop-loxP* fragment was amplified from the Hot-Cre plasmid (Addgene plasmid #24334) and inserted into KpnI/EcoRI sites of the pENTRI1a vector from the Tol2 Kit. Full-length *rxraa* cDNA (ENSDART00000141380.3) was amplified from adult fish hearts cDNA pool using forward primer 5’-atagcggccgcatggaaaccaaacctttctt -3’ and reverse primer 5’-gcggatatcttgtcatttggtgtggagcttct-3’. A clone with the correct sequence of the full-length *rxraa* was confirmed by Sanger sequencing, and the gene was then cloned into the pENTRI1-*loxP-mCherry-stop-loxP* vector. To generate the final construct, p5E-*βactin2*, pENTRI-*loxP-mCherry-stop-loxP-rxraa*, p3E-*EGFP-polyA* and pDestTol2pA were combined together with the Gateway LR clonase II plus enzyme (Thermo Fisher). The final construct was confirmed by gel electrophoresis. 50 ng-100 ng final construct, together with 100 ng transposase mRNA, was then injected into WIK embryos at 1-cell stage. Founder fish (F0) were identified based on mCherry fluorescence. Transgenic fish used for experiments were from F2 and F3 generations.

### Conditional expressional system

*Tg(hsp70l:Cre)*^*zdf13*^, *Tg(kdrl:CreER)*^*fb13*^, *Tg(cmlc2:CreER)*^*pd10*^ and *Tg(tcf21:CreER)*^*pd42*^ were obtained from Drs. Caroline Burns and Geoffrey Burns lab, Dr. Kenneth Poss lab and zebrafish international resource center (ZIRC) (52, 82-84). To activate the heat shock promoter, double-transgenic fish were incubated at 37 °C for 1 hour at 1 dpf. The GFP reporter integrated in *Tg(hsp70l:Cre)*^*zdf13*^ can transiently emit fluorescence about 2-6 hours after incubation and the signal dissipates within the next 48 hours. To activate a tissue-specific promoter, embryos from double-transgenic fish were incubated in 10 µM 4-hydroxytamoxifen (4HT) (Sigma) from 0 dpf to 6 dpf for *Cre-loxP* recombination. E3 water containing 4HT was refreshed every 24 hours. For subsequent experiments on the adult stage, embryos with effective recombination were selected based on eye and body florescence (assessed with a Zeiss microscope) by 6 dpf. Only recombinant-positive embryos were raised and used is the subsequent experiments.

### Compound treatment in zebrafish

For DEAB treatment, compounds were dissolved in dimethyl sulfoxide (DMSO) to create a stock solution. Adult zebrafish were incubated overnight (about 12 hours) in 500 ml system water containing DEAB in a 1L mini tank. We tested DEAB at various concentrations (250 µM, 100 µM, 50 µM, 25 µM, 2.5 µM) and determined that the maximum concentration that did not result in fish death after an overnight incubation was 25 µM. Adult fish were incubated in 25 µM DEAB for 12 hours per day for 2 months. Fresh DEAB water was used daily, and the density of fish was maintained at <5 fish per 500 ml.

For treatment of RXRA agonists at embryonic stages, embryos were incubated in E3 water containing compounds at the desired concentrations. 1-phenyl 2-thiourea (PTU) was used to remove pigmentation. For administration of RXRA agonists to adult fish, bexarotene and isotretinoin were delivered via oral gavage (63) at the desired dose (Supplemental Table 2) on daily basis.

### Adult and embryonic DIC model

For adult zebrafish, DOX was delivered via intraperitoneal injection (20 mg/kg) as previously described (16, 19). For embryonic fish, DOX was delivered from 24 to 72 hpf (53); DOX was dissolved in E3 water containing 100 µM PTU, and the solution was refreshed every 24 hours.

### Quantification of heart pump function via an *ex vivo* system

We employed our recently developed Langendorff-like perfused zebrafish heart technique (44). Briefly, hearts were isolated from tricain-anesthetized fish, cannulated by using 34G ultra-thin catheters through the atrio-ventricular canal visualized with a stereo-microscope Leica M165C (Leica, Germany), paced with an isolated stimulator MyoPacer (Ionoptix; ∼15 V, 10 ms, 2 Hz), and perfused using a peristaltic pump EP-1 EconoPump (BioRad). Images were acquired by using a CMOS camera (MU1403; Amscope; 66 frames per second). All experiments were conducted at room temperature.

End-diastolic and end-systolic volumes (EDV and ESV, respectively) were calculated with the area-length formula:

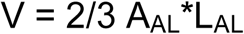

where A_AL_ is area of the base of the ventricle in transverse projection and L_AL_ is long length of the ventricle in longitudinal projection. In order to get 2 perpendicular projections of the ventricle images, we employed a 45° angle aluminum mirror (Thorlabs). The images were analyzed using the ImageJ software (National Institutes of Health); 3 cardiac cycles were analyzed to obtain averaged EDV and ESV values. Ejection fraction was calculated as follows:

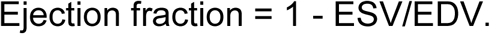

### Quantification of heart pump function via a high-frequency echo system

High-frequency echocardiography was leveraged when quantification of pump function needed to be conducted at multiple time points on the same adult zebrafish heart (Figure 4, 7). All ultrasound movies were documented with a Vevo 3100 high-frequency imaging system (FUJIFILM VisualSonics Inc., Toronto, Canada) equipped with a 50-MHz linear array transducer (MX700). Acoustic gel (Aquasonic 100, Parker Laboratories, Inc) was applied over the surface of the transducer to provide adequate coupling with the tissue interface. Zebrafish were anesthetized with 0.16 mg/mL tricaine, placed ventral side up, and held in place with a soft-sponge stage. The MX700 transducer was positioned above the zebrafish to provide a sagittal imaging plane of the heart. B-mode images were acquired with an imaging field of view of 9.00 mm in the axial direction and 5.73 mm in the lateral direction, a frame rate of 123 Hz, with medium persistence and a transmit focus at the center of the heart. Images were quantified with the VevoLAB workstation. Data were acquired and processed as previously described (51). Ventricular chamber dimensions were measured from B-mode images using the following 2 indices: fractional shortening (FS) = (EDD-ESD)/EDD; fractional area change (FAC) = (EDA-ESA)/EDA. EDD and ESD were the perpendicular distances from the ventricular apex to the ventricular basal line at the end-diastolic and end-systolic stages, respectively; EDA and ESA were defined as the areas of the ventricular chamber at the end-diastolic and end-systolic stages, respectively. For each index, measurements were obtained during 3 to 5 independent cardiac cycles per fish to determine average values.

### Swimming tunnel assay

Swimming challenge was conducted on adult zebrafish using a swim tunnel respirometer (Mini Swim 170, Loligo Systems, Tjele, Denmark), with protocol that was previously reported (85, 86). Critical swimming speed (Ucrit) was defined as the maximum water speed that fish were able to swim against while maintaining their position. Fish were fasted for 24 hours for synchronization before the test. Five to 10 fish were loaded into the swim tunnel and acclimated in a lower speed of 9 cm/s (200 rpm) of flowing water for 20 minutes. Water speed was then increased in steps of 8.66 cm/s (100 rbm) (Uii) at 150-second intervals (Tii) until all fish were exhausted and failed to resume swimming from a downstream screen. The highest water speed (Ui) against which fish were able to complete the 150-second swim test, and the swimming duration time (Ti) in the next 150-second period, were recorded for each fish. Critical swimming speed was calculated with the following formula (87):

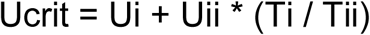

and normalized to fish body length for intergroup comparisons.

### Human coronary artery endothelial cell culture

Primary human coronary artery endothelial cell (HCAEC) were purchased from ATCC (PCS-100-020) and cultured in vascular cell base medium (PCS-100-030) containing endothelial cell growth kit (PCS-100-041) at 37 °C. HCAEC at the fourth or fifth passage were used for DOX treatment. For pretreatment of RXRA agonists, HCAEC were treated with isotretinoin or bexarotene for 24 hours and then subjected to DOX stress. For cotreatment of RXRA agonists, culture medium containing DOX and isotretinoin or bexarotene was added together. All cells were harvested 24 hours after DOX treatment.

### Real time quantitative PCR

For fish embryo studies, 20 embryos were pooled for RNA extraction. For adult fish studies, 3 freshly dissected tissues were pooled for RNA extraction. For primary cell culture, cells in a 60-mm plate were harvested as a single biological replicate. RNA was extracted using Trizol (Biorad) following manufacturer instructions. cDNA was synthesized by using Superscript III First-Strand Synthesis System (Invitrogen) and 100-500 ng of RNA. Real-time RT-PCR assays were performed in 96-well optical plates (ThermoFisher) using an Applied Biosystem VAii 7 System (ThermoFisher). Primers are listed in Supplemental Table 3 and 4. Levels of gene expression were normalized to either *glyceraldehyde 3-phosphate dehydrogenase* (*gapdh)* or *actin beta 2* (*actb2*).

### Western blotting

Zebrafish embryos at 5 dpf or freshly dissected adult ventricular tissues were collected in 1.5 mL safe-lock tubes (Eppendorf) and mechanically homogenized (Blender tissue homogenizer; Next Advance, Inc) in RIPA lysis buffer (Sigma) containing proteinase inhibitor, 1 mM phenylmethylsulphonyl fluoride (PMSF), and stainless steel beads. Standard protocols of western blotting were followed. For each sample, >20 µg of total protein was loaded on a sodium dodecyl sulfate (SDS) polyacrylamide gel. The following primary antibodies were used: anti-EGFP (1:2,000) (Santa Cruz), anti-mCherry (1:1,000) (Novus Biologicals), anti-cleaved caspase 3 (1:1,000) (Cell Signaling Technology). Anti-actin (1:2,000) (Sigma) was used as an internal control.

### Immunofluorescence

Embryonic hearts were dissected by following a published protocol (88). Dissected embryonic hearts were fixed in 4% paraformaldehyde (PFA) before imaging. Adult cardiac tissues were freshly dissected and placed in molds with frozen-section medium on dry ice. Frozen samples were transferred to −80°C overnight and subsequently sliced into 10 µm sections with a cryostat (Leica CM3050S). For endogenous mRFP, EGFP and mCherry, images were obtained within 24 hours after dissection of embryonic and adult hearts. For immunostaining, sections of adult cardiac tissues were fixed in 4% PFA and then subject to a standard protocol. Primary antibodies against Mef2 (1:200) (Santa Cruz) and α-actinin (1:100) (Sigma) were used. To quantify apoptotic cells, terminal deoxynucleotidyl transferase dUTP nick and labeling (TUNEL) staining was conducted using an In-situ Cell Death Detection kit (Roche) following manufacturer instruction. All images were captured using a Zeiss Axioplan II microscope equipped with ApoTome 2 and ZEN software (Carl Zeiss).

### Statistics

No sample sizes were calculated before performing the experiments, and no animals were excluded for analyses. Wild type sibling zebrafish were used as a control for mutants or transgenic lines whenever possible. For homozygote mutant lines, age-matched wild type zebrafish were used as controls. For survival analysis, experiments on >3 independent groups of fish were conducted. Log-rank tests are used to compare combined data from all experimental replicates. For dot-plot graphs, values are displayed as mean ± standard deviation. Unpaired 2-tailed student *t* tests were used to compare 2 groups; 1-way ANOVA followed by posthoc turkey tests were used for comparing 3 and more groups, and equal variance was assumed. *P* values less than 0.05 were considered statistically significant. All statistical analyses were conducted by using JMP10 (SAS Institude Inc) and Prism 6 (Graphpad) software.

## Supporting information

## Author contributions

XM and XX conceived the project, designed the experiments, interpreted the data, and drafted the original manuscript. XM performed experiments, collected the data and generated figures. YD contributed to the generation of transgenic animals. HZ and AD performed *ex vivo* cardiac pump assays. QQ and ML evaluated the compounds treatment and performed echocardiography on adult zebrafish. MK performed cell culture-related experiments. YW evaluated the compounds treatment in embryonic zebrafish. XX supervised the research. JH, SCE, TKH, and XL made critical intellectual contributions and edited the manuscript.

### Acknowledgements

This work was supported in part by Scientist Development Grants from the American Heart Association (14SDG18160021 to YD, 15SDG25080264 to MK), the US NIH R01 grants HL 81753, HL 107304, and HL111437 to XX, and the Mayo Foundation to XX. This work was also made possible by CTSA grant UL1TR002377 from the National Center for Advancing Translational Sciences (NCATS), a component of the National Institutes of Health (NIH).

We thank Kashia Stragey and Beninio Gore for maintaining the zebrafish facility at Mayo Clinic. We thank Caroline Burns, Geoffrey Burns, Kenneth Poss, and ZIRC for generously sharing transgenic animals; Michael Underhill for generously sharing RA-related plasmids; and Yue Yu on generously assisting with R coding.

