## Supplementary material for "Retinoid X receptor alpha is a spatiotemporally-specific therapeutic target for doxorubicin-induced cardiomyopathy in adult zebrafish"

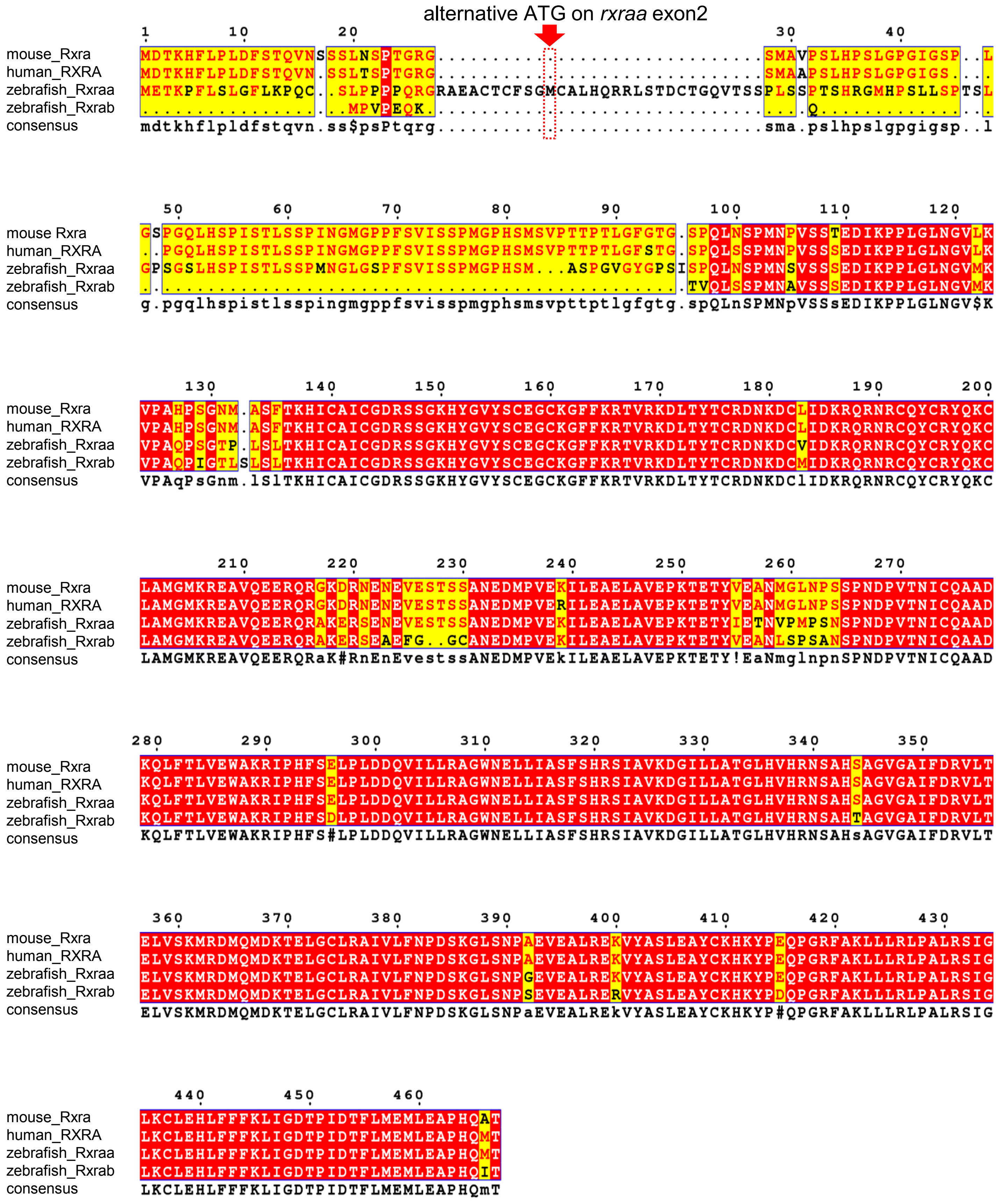


**Supplemental Figure 1.**

**Supplemental Figure 1. RXRA is conserved across species**
Comparison of amino acid sequence of RXRA orthologs among human, mouse and zebrafish. Consensus regions among all species were labeled in red. One predicted alternative ATG start codon after RP2 insertional locus was noted.

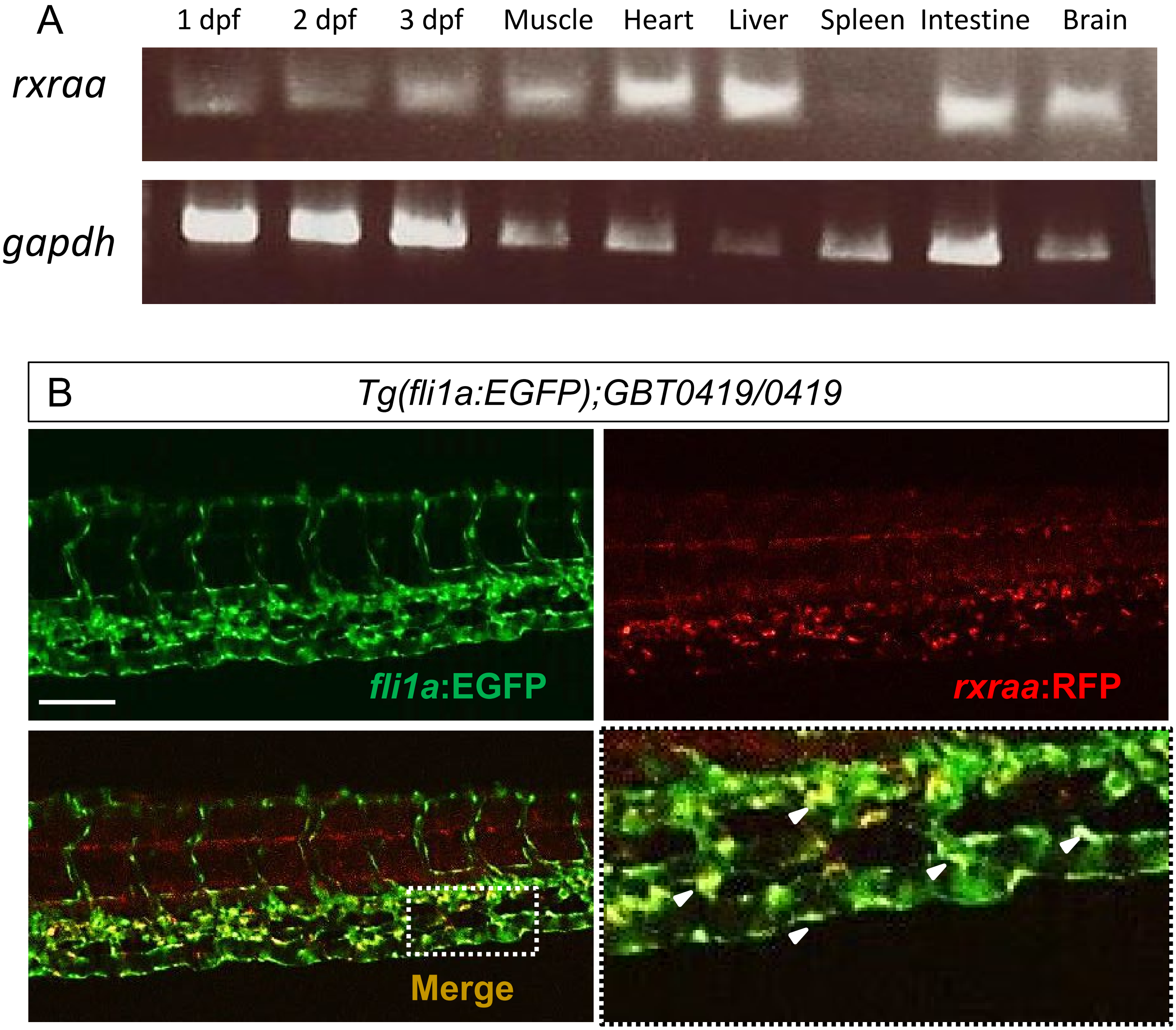


**Supplemental Figure 2.**

**Supplemental Figure 2. Expression of *rxraa* in zebrafish**

**A,** Expression of *rxraa* in embryos and different adult tissues via semi-quantitative RT-PCR. Whole embryos were collected at the indicated stages. Adult tissues were dissected from 3-month-old fish. dpf, days post fertilization. *gapdh*, house-keeping control. **B,** Florescence images of *Tg(fli1a:EGFP);GBT0419/0419* embryo (trunk region) at 2 dpf suggests endothelial expression of *rxraa*. Green fluorescence labels endothelial cells, while red fluorescence labels endogenous *rxraa*. Higher magnification image was shown for the boxed area of the merge channel. Arrows represent overlaps between green and red fluorescence. Scale bar=200 μm.

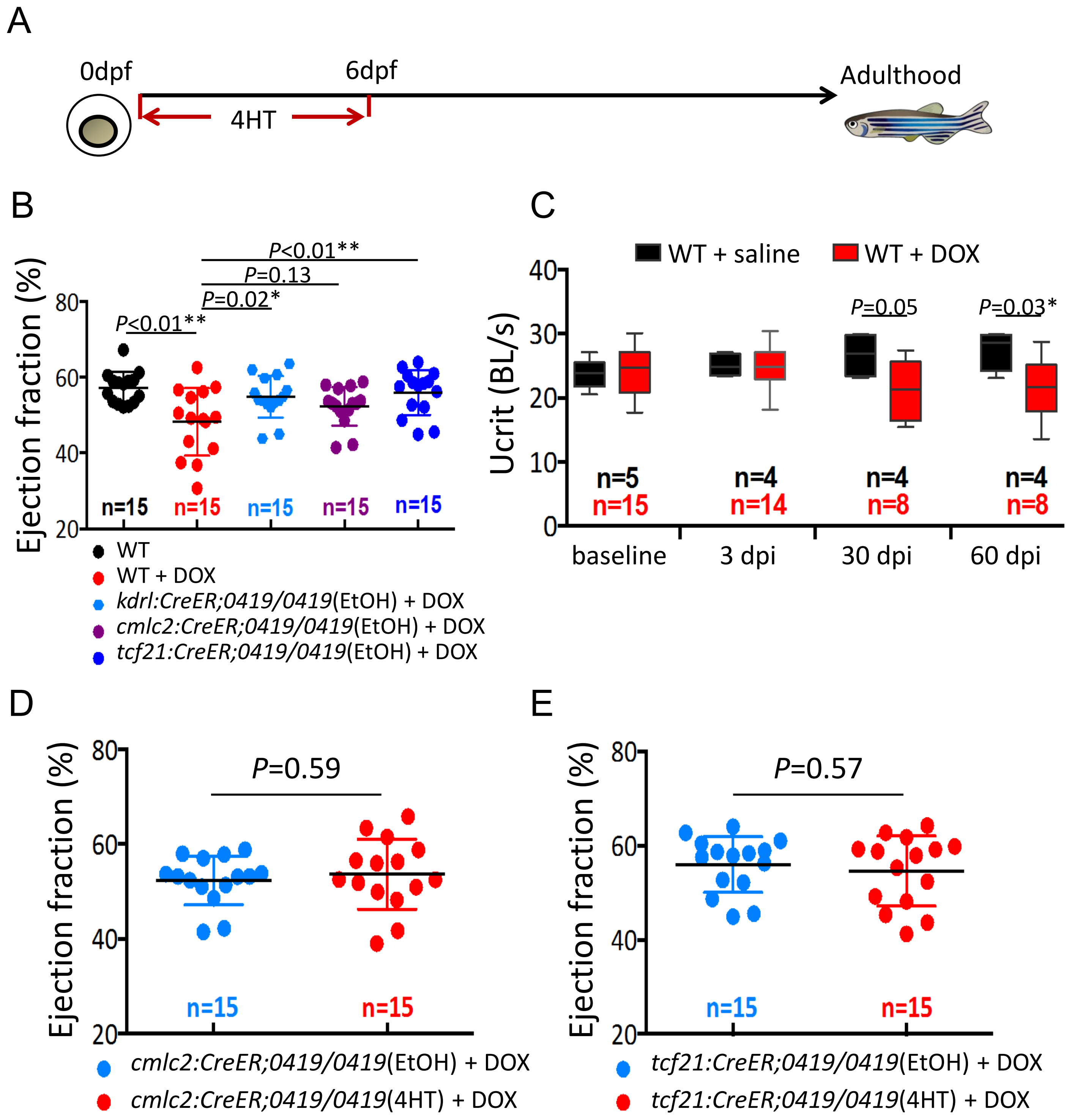


**Supplemental Figure 3.**

**Supplemental Figure 3. Endothelial RP2 reversion in *GBT0419/0419* attenuates its cardioprotective effects against DIC**

**A,** Schematics of the protocol for generating conditional transgenic fish. Transgenic fish were incubate in 10 μM 4-HT in E3 water from 0 dpf (1 cell stage) to 6 dpf. Fish were then transferred to circulating system to grow to adulthood. **B,** Ventricular ejection fraction of WT and *GBT0419/0419* with different *CreER* backgrounds (without 4HT induction) after DOX stress. EtOH, ethonal. **C,** Swimming capacity of adult fish after injected with a single bolus of DOX. Ucrit, critical swimming speed; BL, body length; dpi, days post DOX injection. **D, E,** Ventricular ejection fraction of *GBT0419/0419* adult fish with myocardial **(D)** or epicardial **(E)** RP2 reversion after DOX stress. Each dot represents ejection fraction measurement of a single ventricle. Error bars indicate standard deviation. **P*<0.05, ***P*<0.01, 1-way ANOVA followed by posthoc Tukey test in **(B)**, unpaired student *t* test in **(C)**, **(D)** and **(E)**.

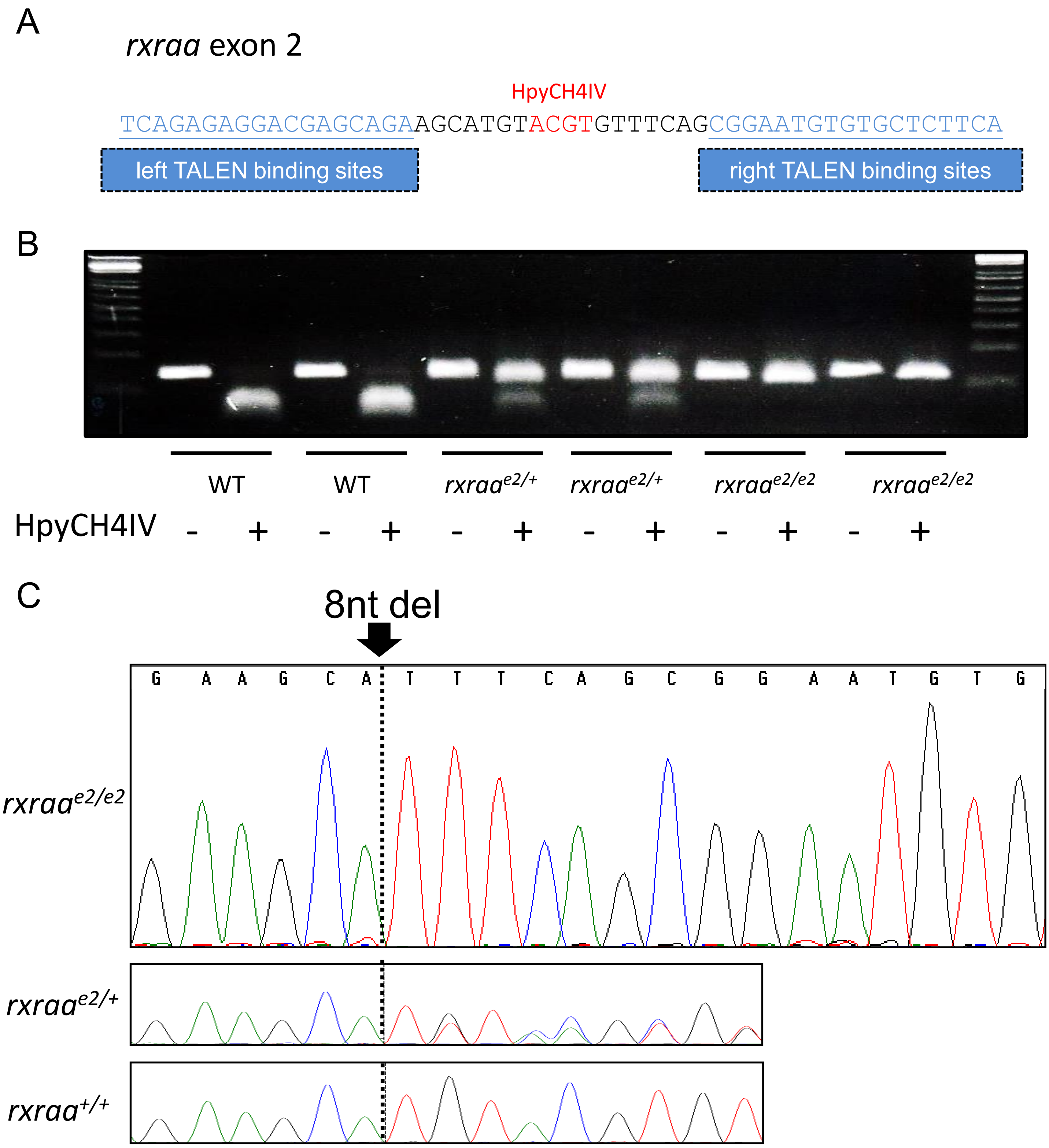


**Supplemental Figure 4.**

**Supplemental Figure 4. Generation of a *rxraa* zebrafish mutant by TALEN**

**A,** Design of TALEN targeting sites on *rxraa* exon 2. TALEN binding sequences are shown in blue. A HpyCH4IV digestion site (red) was harnessed for genotyping to identify successful mutagenesis. **B,** A representative genotyping gel for $rxraa^{e2/+}$ and $rxraa^{e2/e2}$. Successful mutagenesis is indicated by depletion of the HpyCH4IV digestion site. **C,** Sanger sequencing indicated a 8 nucleotides deletion in $rxraa^{e2}$.

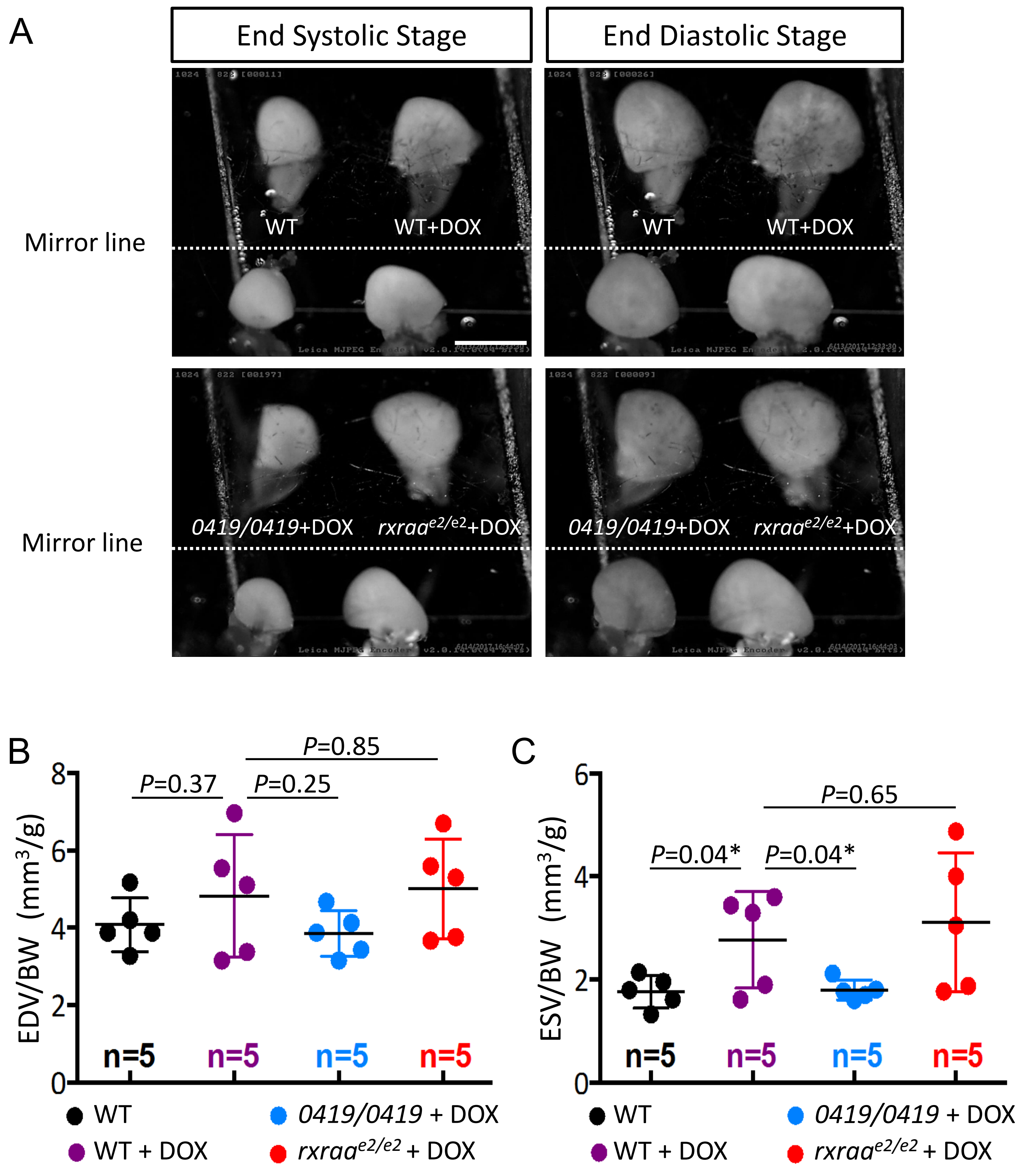


**Supplemental Figure 5.**

**Supplemental Figure 5. *GBT0419/0419* rescues increased end-systolic volume in DIC model**

**A,** Representative images of adult zebrafish hearts at end systolic stage and end diastolic stage of a beating cycle in *ex vivo* system. Images on two sides of mirror represent the same heart with 90° reflections. Scale bar=1 mm. **B, C,** Quantification of end systolic volume **(B)** and end diastolic volume **(C)** of pumping ventricles. Each dots represents a single adult ventricle. Indices were normalized to body weight. ESV, end systolic volume; EDV, end diastolic volume; BW, body weight. Error bars indicate standard deviation. **P*<0.05, 1-way ANOVA followed by posthoc Tukey test.

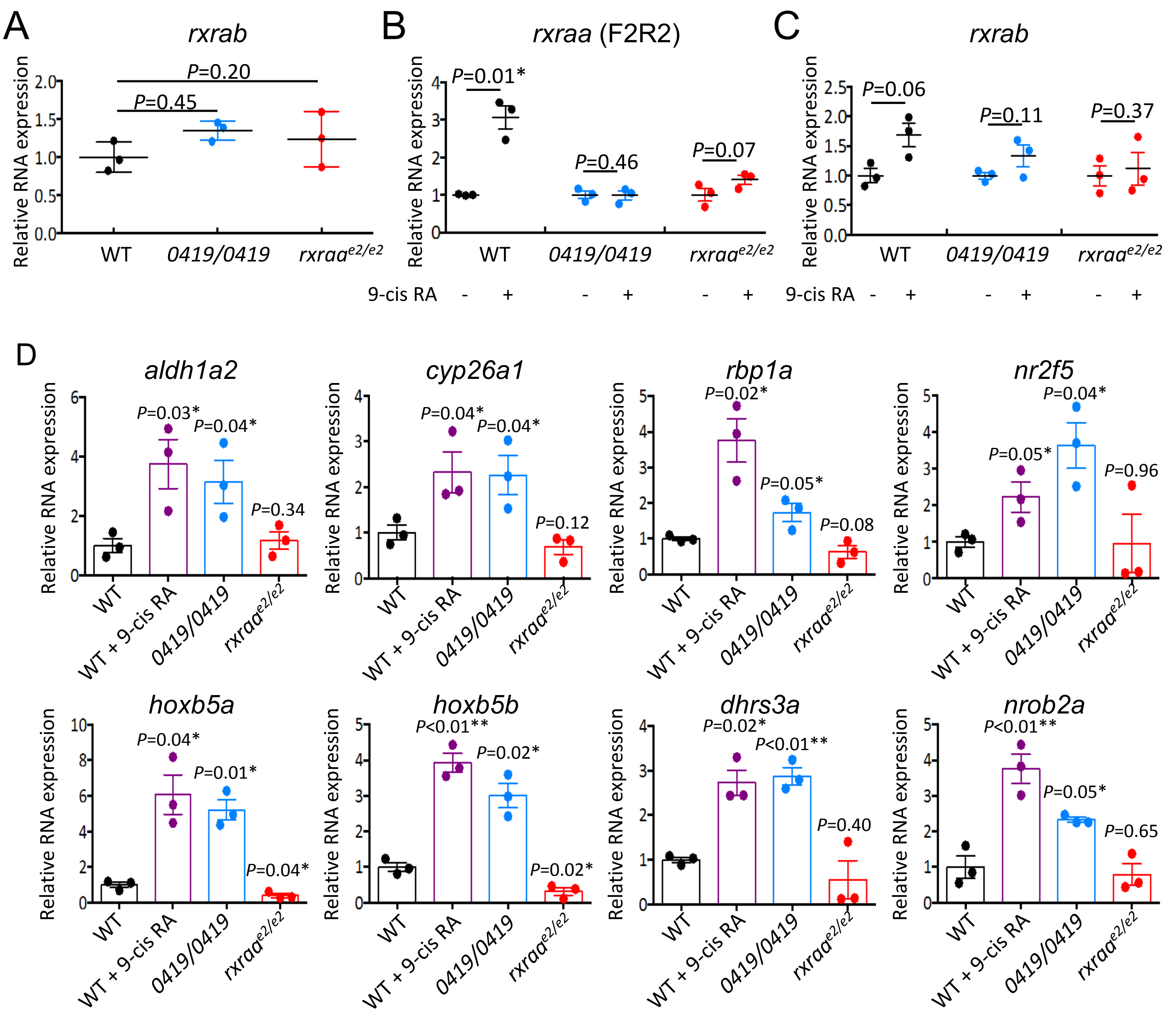


**Supplemental Figure 6.**

**Supplemental Figure 6. Hyper-activation of RA signaling in *GBT0419/0419***

**A,** Relative mRNA expression of *rxrab* in WT, *GBT0419/0419* and *rxraa*$^{e2/e2}$. **B,** Relative mRNA expression of *rxraa* upon 9-cis RA treatment. **C,** Relative mRNA expression of *rxrab* upon 9-cis RA treatment. 9-cis RA was adminstered from 24 hpf to 48 hpf. **D,** mRNA expressions of RA signaling-responsive gene targets in *GBT0419/0419* and *rxraa*$^{e2/e2}$. As a positive control, we noted activated RA signaling-responsive gene targets upon treatment of 9-cis RA. These data were also shown as a heatmap in **Figure 4D**. Error bars show standard deviation. *$P<0.05$, **$P<0.01$, 1-way ANOVA followed by posthoc Tukey test in **(A)** and **(D)**; shown statistics in **(D)** represents comparison between corresponding group with WT group; unpaired student *t* test in **(B)** and **(C)**.

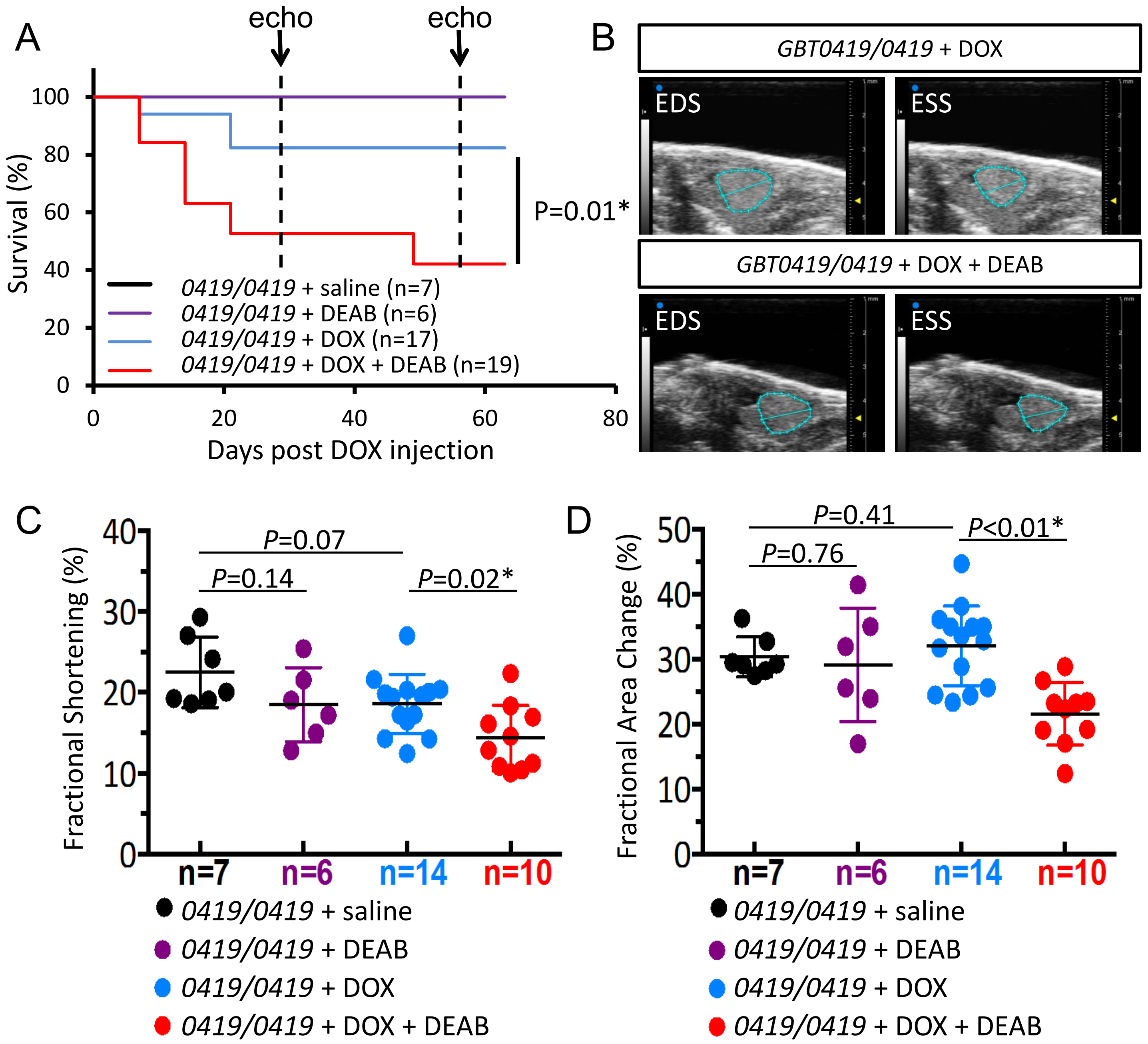


**Supplemental Figure 7.**

**Supplemental Figure 7. Inhibition of RA signaling reduced survival and cardiac function in *GBT0419/0419* upon DOX stress**

**A,** Shown are Kaplan-Meier survival curves of DOX stressed *GBT0419/0419* adult fish with or without DEAB treatment. Echocardiography was performed at 4 wpi and 8 wpi. The black curve overlaps with the purple curve. **B,** Representative echocardiography images of DOX stressed *GBT0419/0419* adult fish at 4 wpi. EDS, end systolic stage; ESS, end systolic stage. **C,** Ventricular fractional shortening of *GBT0419/0419* adult fish, with DEAB treatment at 4 wpi after DOX stress. **D,** Ventricular fractional area change of *GBT0419/0419* adult fish, with DEAB treatment, at 4 wpi after DOX stress. Error bars show standard deviation. *$P<0.05$, **$P<0.01$, 1-way ANOVA followed by posthoc Tukey test in **(C)** and **(D)**, Log-rank test in **(A)**. Saline were used as sham treatment.

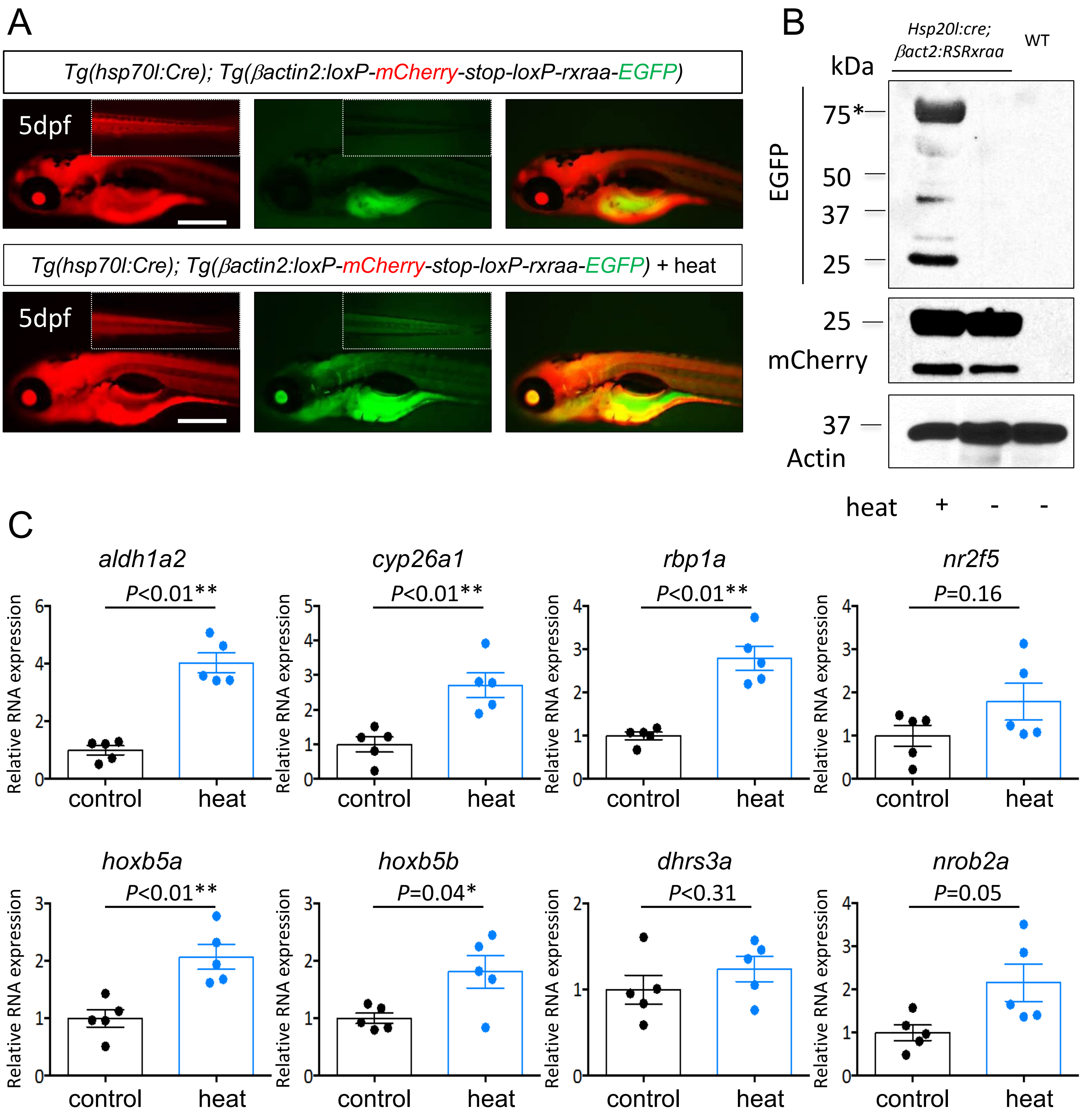


**Supplemental Figure 8.**

**Supplemental Figure 8. Ectopic expression of *rxraa* activates RA signaling *in vivo***

**A,** Images of 5 dpf embryos of *Tg(hsp70l:cre); Tg(βact2:RSRxraa)* after heat shock. Insets show tail region of the same embryo on the same scale. Scale bar=500 μm. **B,** Shown are Western blot to confirm Rxraa-EGFP fusion protein induction, which can be detected by an anti-EGFP antibody, after transgenic embryos were heat shocked. Predicted size of fusion protein is 75 KDa (Rxraa, 50KDa; EGFP, 25KDa) and is indicated by an asterisk. mCherry was detected in all embryos with the *Tg(βact2:RSRxraa)* transgene. Actin was used as an internal control. **C,** mRNA expression of RA signaling-responsive gene targets in *Tg(hsp70l:cre); Tg(βact2:RSRxraa)* with or without heat shock. control, no heat. Error bars indicate standard deviation. **P*<0.05, ***P*<0.01, unpaired student *t* test was used.

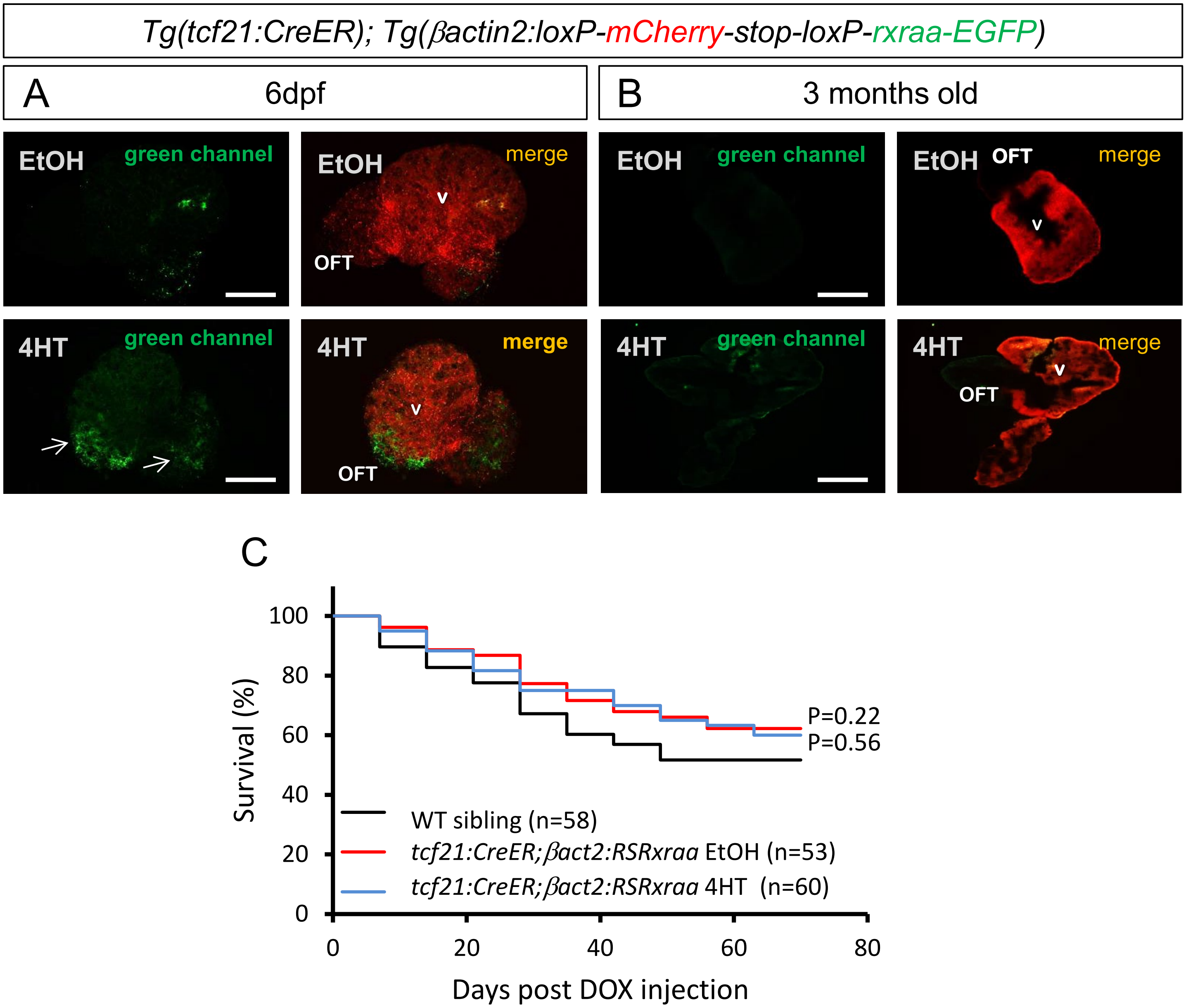


**Supplemental Figure 9.**

**Supplemental Figure 9. Induction of Rxraa-fusion protein in *tcf21*⁺ epicardial cells**

**A,** Representative florescence images of 6 dpf *Tg(tcf21:CreER); Tg(βact2:RSRxraa)* dissected hearts with or without *Cre-loxP* recombination. Arrows indicate induced green florescence signal in the ventricle and outflow tract. Scale bar=100 μm. **B,** Representative florescence images of 3-month-old *Tg(tcf21:CreER); Tg(βact2:RSRxraa)* dissected hearts with or without *Cre-loxP* recombination. Scale bar = 500 μm. In **(A)** and **(B)**, Fish were treated with 4HT from 0 dpf to 6 dpf. V, ventricle; OFT, outflow tract; EtOH, ethonal. **C,** Show are Kaplan-Meier survival curves for DOX stressed adult fish with epicardial-specific induction of ectopic *rxraa*. Log-rank test was performed, shown statistics were comparisons to WT sibling group.

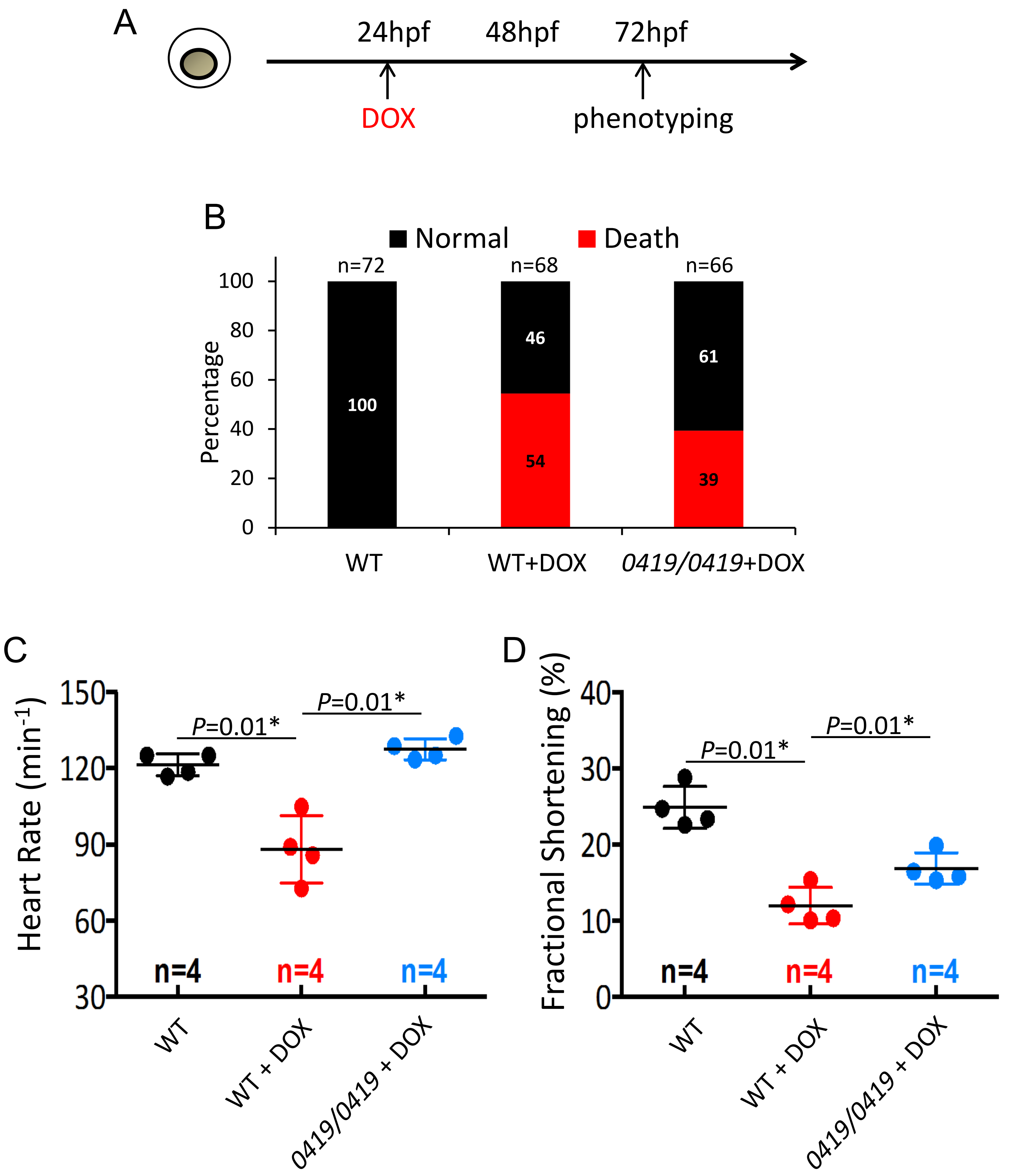


**Supplemental Figure 10.**

**Supplemental Figure 10.** ***GBT0419/0419*** **preserve cardiac function in a zebrafish embryonic DIC model**

**A,** The design of an embryonic zebrafish DIC model. hpf, hours post fertilization. **B,** Survival percentages of embryos after DOX treatment at 72 hpf. Numbers on the plots represent proportion of either dead or living embryos. **C,** Heart rate of embryos after DOX treatment. **D,** Ventricular fractional shortening after DOX treatment. 100 μM DOX was used. *$P<0.05$, **$P<0.01$, 1-way ANOVA followed by posthoc Tukey test.

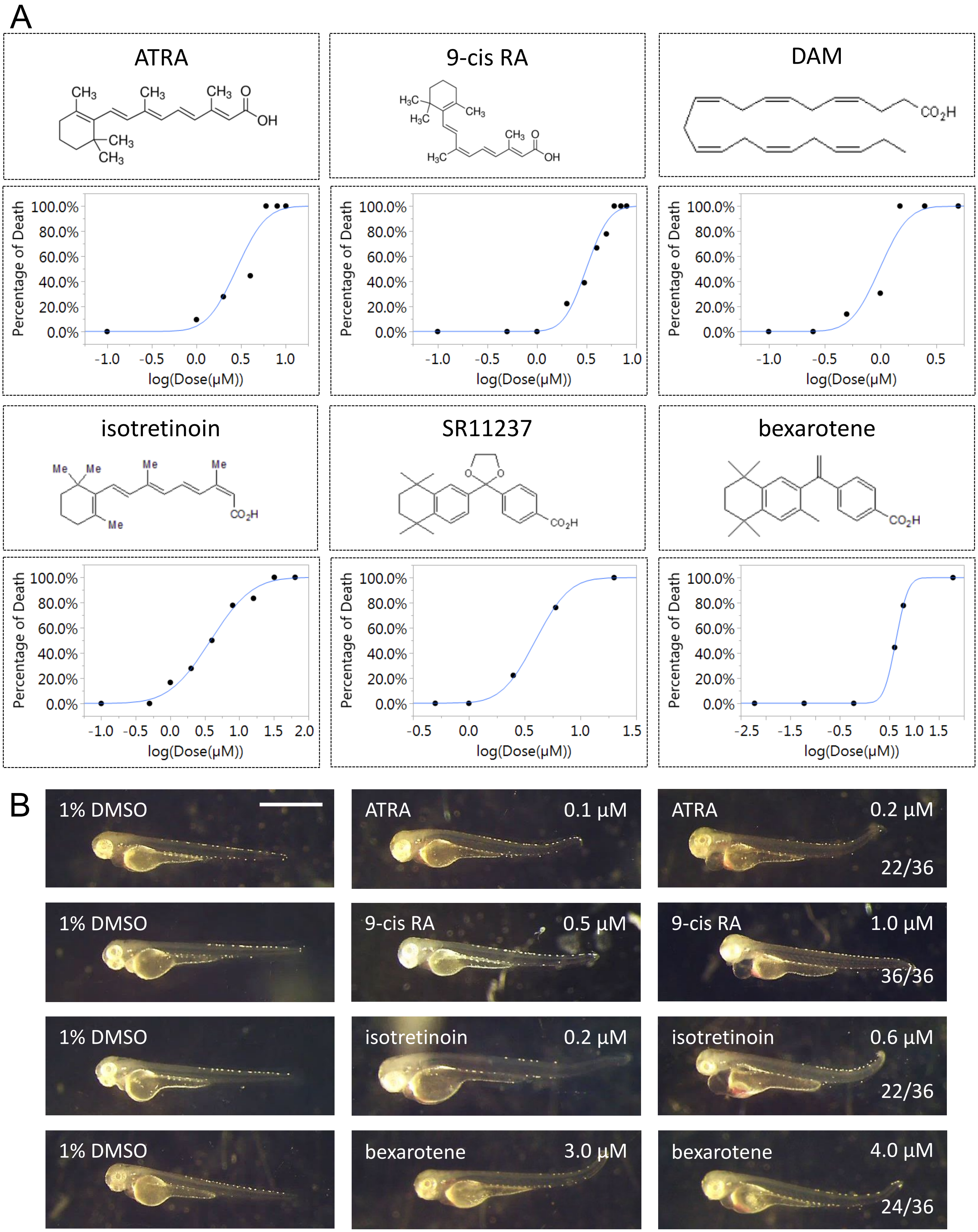


**Supplemental Figure 11.**

**Supplemental Figure 11. LD50 and impacts of high-dose RXRA agonists on zebrafish embryos**

**A,** The median lethal dose (LD50) of each RXRA agonists were experimentally determined. For each RXRA agonist, their chemical structures were shown in the upper panels, while percentage of embryo lethality at a series of different doses were shown in the lower panels. n>18 embryo were tested at each dose. **B,** Representative cardiac teratogenic phenotypes caused by RXRA agonists. Left, control embryos treated with 1% DMSO. Middle, embryos show no cardiac phenotypes were noted if the dose is less than the listed dose. Right, some embryos exhibit cardiac phenotypes when treated at listed high dose. Numbers at the bottom right indicate proportion of animals with phenotypes.

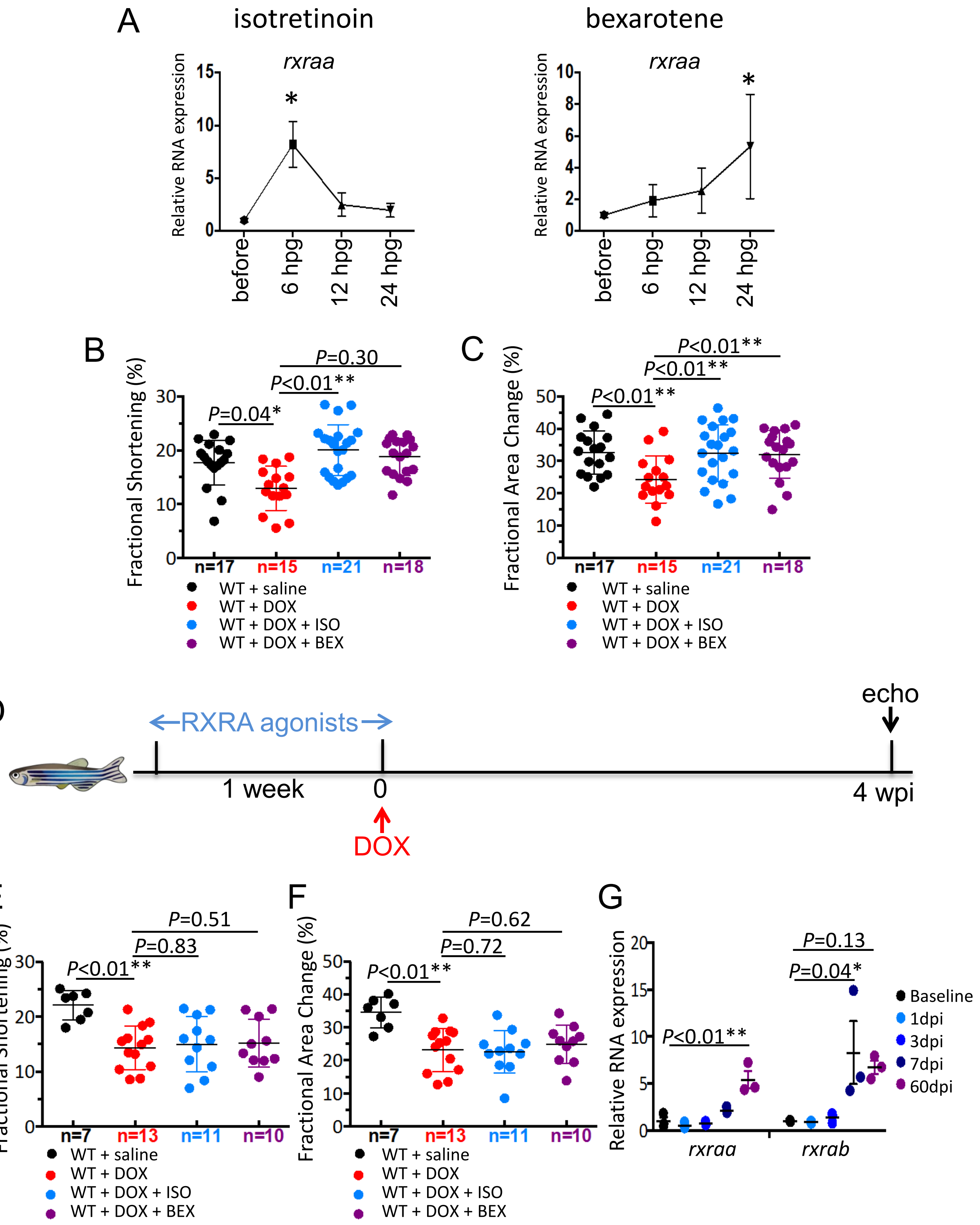


**Supplemental Figure 12.**

**Supplemental Figure 12. Stage-dependent treatment of RXRA agonists on adult DIC**

**A,** Responses of RA signaling in the heart to one-time oral gavage to 2 RXRA agonists at clinical relevant doses shown in **Supplemental Table 2.** Shown are real time RT-PCR for the expression of *rxraa* and *aldh1a2* at different time points after gavage. before, before gavage; hpg, hours post gavage. **B,** Fractional shortening of adult ventricle at 4 wpi in fish treated with ISO and BEX during the early DIC stage. **C,** Fractional area change of adult ventricle at 4 wpi in fish treated with ISO and BEX during the early DIC stage. **D,** Schematic for the design of pre-treatment of RXRA agonists in the adult fish DIC model. RXRA agonists were administrated daily by oral gavage for one week before a single bolus of DOX was injected. **E,** Fractional shortening of adult ventricle at 4 wpi in fish pre-pretreated with ISO or BEX. **F,** Fractional area change of adult ventricle at 4 wpi in fish pre-pretreated with ISO or BEX. **G,** Dynamics of mRNA expression of cardiac *rxraa* and *rxrab* after DOX stress. >90 WT fish were stressed with DOX, and RNA was extracted from a pool of 3 dissected ventricles at baseline (before DOX injection), 1 dpi, 3 dpi, 7 dpi and 60 dpi, respectively. Samples were collected in triplicates at each time point. dpi, days post DOX injection; wpi, weeks post DOX injection; ISO, isotretinoin; BEX, bexarotene; $^*P<0.05$, $^{**}P<0.01$, 1-way ANOVA followed by posthoc Tukey test.

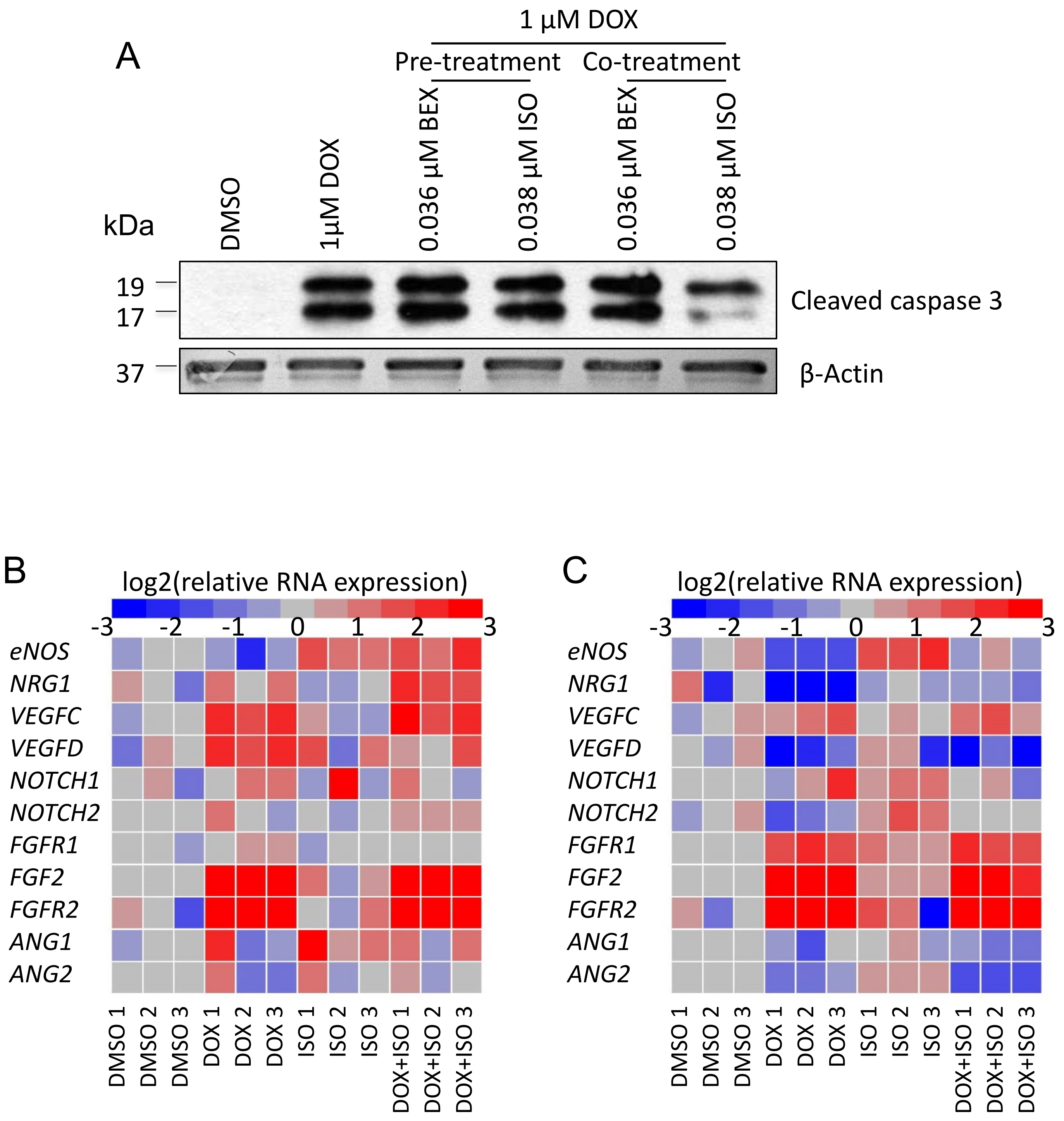


**Supplemental Figure 13.**

**Supplemental Figure 13. Treatment of RXRA agonists in human coronary artery endothelial cell**

**A,** Show are western blot to measure cleavage caspase 3 as an apoptosis index. βactin is used as an internal control. **B, C,** Heatmaps showing quantitative RT-PCR results of key genes expression in endothelial paracrine signaling pathways. *gapdh* is used as an internal control. Values are shown as $\log_2$ transformation of relative expression to mean values of control group (DMSO). **D, E,** mRNA expression of endothelial *nitric oxide synthase* (*eNOS*). DOX, doxorubicin; ISO, isotretinoin; BEX, bexarotene. *$P$<0.05, **$P$<0.01, 1-way ANOVA followed by posthoc Tukey test.

**Supplemental Table 1. Cardiac expression of zebrafish orthologs for *RXRA***

| Gene | Ensembl ID | Chromosome | embryonic heart RPKM | adult heart RPKM |
|---|---|---|---|---|
| *rxraa* | ENSDARG00000057737 | chr21 | 0.85±0.03 | 1.17±0.07 |
| *rxrab* | ENSDARG00000035127 | chr5 | 0.25±0.02 | 0.29±0.11 |

Data were extracted from published RNA sequencing data (GEO, #GSE85416).
RPKM, reads per kilobase million.

**Supplemental Table 2. Dose conversion of isotretinoin and bexarotene from human to fish**

| Compound | Human Dose (mg/m²/day) | BSA of a 60kg patient (m²) | Dose for a 60kg patient (mg/day) |
|---|---|---|---|
| isotretinoin | 74 | 1.6 | 118 |
| bexarotene | 300 | 1.6 | 480 |
| **Compound** | **Fish Dose (mg/m²/day)** | **BSA of a 0.3g zebrafish (cm²)** | **Dose for a 0.3g zebrafish (µg/day)** |
| isotretinoin | 74 | 3.2 | 24 |
| bexarotene | 300 | 3.2 | 97 |

1. Human doses were obtained according to FDA guidelines: FDA reference ID3820564 and ID2946940.
2. Dosage conversion from human to adult zebrafish were calculated based on body surface area (BSA). BSA of adult zebrafish were measured experimentally.

**Supplemental Table 3. Sequences for quantitative PCR primers in zebrafish**

| Zebrafish Gene | Ensembl ID | Forward | Reverse |
|---|---|---|---|
| *actb2* | ENSDARG00000037870 | ggtatcgtgatggactctgg | tctcctgctcaaagtcaagg |
| *aldh1a2* | ENSDARG00000053493 | gtttgaacagtagcttccccc | gctaccctggagtctctgga |
| *cyp26a1* | ENSDARG00000033999 | agagatgaagcggctgatgt | tcttcttgctgctgtcgatg |
| *dhrs3a* | ENSDARG00000044982 | caacactgccttcgttgtc | gtgttcatacaggtgtagcttc |
| *gapdh* | ENSDARG00000043457 | ccacccatggaaagtacaag | ctctctttgcaccaccctta |
| *hoxb5a* | ENSDARG00000013057 | tcccttggatgaggaaactac | atttgacgctctgagaggc |
| *hoxb5b* | ENSDARG00000054030 | aactccacagatattcccctg | cgtctggtagcgagtgtaag |
| *nppa* | ENSDARG00000052960 | gatgtacaagcgcacacgtt | tctgatgcctcttctgttgc |
| *nppb* | ENSDARG00000052958 | catgggtgttttaaagtttctcc | cttcaatatttgccgcctttac |
| *nr2f5* | ENSDARG00000033172 | aaagcccttcaggtggacac | agggaagccgtaacaacagg |
| *nr0b2a* | ENSDARG00000044685 | cctgcatcaacaagttctgg | agcaaaacgtcctccatcc |
| *rxraa* (F1R1) | ENSDARG00000057737 | atccgtcttgcagtgtcatcagagagag | ctaatgaagttggactcagcagtgac |
| *rxraa* (F2R2) | ENSDARG00000057737 | gcatctcctggagtgggtta | tggacggcttctctcttcat |
| *rxrab* | ENSDARG00000035127 | ccatggggatgaagagagaa | ttcacagctatggagcgatg |

**Supplemental Table 4. Sequences for quantitative PCR primers in primary cell culture.**

| Human Gene | reference | Forward | Reverse |
|---|---|---|---|
| *GAPDH* | (Chang, Fu et al. 2011) | cagcaagagcacaagaggaagaga | ttgatggtacatgacaaggtgcgg |
| *eNOS* | (Lapointe, Roy et al. 2006) | ccttccgctaccagccaga | cagagatcttcactgcattggcta |
| *NRG1* | (Wang, Zhou et al. 2017) | cttcggtcagaacggagcaa | acagtcgtggagtgatgggc |
| *VEGFC* | (Leclers, Durand et al. 2006) | cacgagctacctcagcaaga | gctgcctgacactgtggta |
| *VEGFD* | (Leclers, Durand et al. 2006) | cctgaagaagatcgctgttc | gagagctggttcctggagat |
| *NOTCH1* | (Ishida, Hijioka et al. 2013) | caatgtggatgccgcagttgtg | cagcaccttggcggtctcgta |
| *NOTCH2* | (Baeten and Lilly 2015) | acagttgtgtctgctcaccaggat | gcggaaaccattcacaccgttgat |
| *FGFR1* | (Seo, Jeong et al. 2016) | tgagaagaagacaccaagtgga | ttgtggcgctttcaagacta |
| *FGF2* | (Igarashi, Okamoto et al. 2016) | agcggctgtactgcaaaaac | gcttgaagttgtagcttgatgtg |
| *FGFR2* | (Seo, Jeong et al. 2016) | tctgcgtttggagttgctc | gctgctgctgcagtcactt |
| *ANG1* | (Ward, Haninec et al. 2004) | tgggaagggaacgagcctatt | aatcatcatagttgtggaacgtaa |
| *ANG2* | (Hegen, Koidl et al. 2004) | cagttcttcaaaagcagcaacatg | gatccggatgtttagaaatctgctcgt |
